# Drug-induced colorectal cancer persister cells show increased mutation rate

**DOI:** 10.1101/2021.05.17.444478

**Authors:** Mariangela Russo, Simone Pompei, Alberto Sogari, Mattia Corigliano, Giovanni Crisafulli, Andrea Bertotti, Marco Gherardi, Federica Di Nicolantonio, Alberto Bardelli, Marco Cosentino Lagomarsino

**Affiliations:** Department of Oncology, University of Torino, Candiolo (TO), Italy; Candiolo Cancer Institute, FPO–IRCCS, Candiolo (TO), Italy; IFOM Foundation, FIRC Institute of Molecular Oncology, Milan, Italy; Dipartimento di Fisica, Università degli Studi di Milano, and I.N.F.N, via Celoria 16 Milano, Italy

## Abstract

Compelling evidence shows that cancer persister cells limit the efficacy of targeted therapies. However, it is unclear whether persister cells are induced by anticancer drugs, and if their mutation rate quantitatively increases during treatment. Here, combining experimental characterization and mathematical modeling, we show that, in colorectal cancer, persisters are induced by drug treatment and show a 7- to 50-fold increase of mutation rate when exposed to clinically approved targeted therapies. These findings reveal that treatment may influence persistence and mutability in cancer cells and pinpoints new strategies to restrict tumor recurrence.

## Main Text

When cancer patients are treated with targeted agents such as EGFR or BRAF inhibitors, after an initial response, tumor relapse is often observed^1,2^. A widely accepted view is that cells harboring drug resistant mutations are already present in the tumor mass before treatment initiation and subsequently repopulate the lesion in a matter of months^3,4^ However, emergence of drug resistance after prolonged response and disease stabilization is also frequently reported^5,6^. Since cells harboring pre-existing mutations can proliferate in the presence of the drug, it is unclear why they would require such a long time (sometimes years) to become clinically detectable. Indeed, when cancer cells are exposed to lethal doses of targeted therapies, the emergence of a sub-population of drug-tolerant persister cells is often observed^7–11^, limiting tumor eradication^12^.

Although persister cells have been described across multiple cancer types in response to different therapies^7,8,13–15^, their phenotype has not been fully characterized. Persister cells are able to survive lethal doses of therapeutic agents and represent a reservoir from which heterogenous mechanisms of drug resistance arise^7,8,16^. Differently from genetically resistant cells, persisters tolerate drug pressure through reversible, non-genetical, non-inheritable mechanisms of resistance^7,12,17^. However, it is unclear if persister cells enter a fully quiescent state or slowly progress through the cell cycle. It is also unknown if the persister phenotype is drug-induced, at least to some extent, or it entirely preexists exposure to therapies (so that drugs would simply select a pre-existing sub-population). Additionally, the population dynamics governing persisters evolution to resistance have only been partially elucidated^10^.

We and others previously reported that cancer cells that survive to lethal doses of anti-cancer agents have the ability to resume growth and re-acquire drug sensitivity upon drug withdrawal, as expected from persisters, and to generate new sub-population of residual persister cells upon further exposure to the same treatment^7,12,18^. However, to distinguish persistence from tolerance, in bacteria exposed to antibiotic treatment, the presence of persisters is quantitatively defined by a biphasic pattern of death curve^19^. Such stringent definitions are often lacking in the context of cancer^12^.

Using colorectal cancer (CRC) as a model system, we previously reported that exposure of tumor cells to targeted therapies induces DNA damage and activation of a stress response characterized by impairment of DNA mismatch repair (MMR) and homologous recombination (HR) proficiency and a switch from high- to low-fidelity DNA replication mediated by error-prone DNA polymerases^18^. This phenotype has been recently confirmed in multiple cancer types in response to different targeted therapies^20^. Drug treatment leads to error-prone DNA replication in cancer cells, supporting the hypothesis that the mutation rate of persister cancer cells could increase during therapy-induced stress^21^. However, mutation rates of cancer cells during treatment with targeted agents have not been quantitatively assessed.

Measuring mutational processes by DNA sequencing is challenging, owing to tumor heterogeneity and to the difficulties of tracking lineages^20^. A complementary strategy is represented by the so-called “fluctuation test”, originally developed by Luria and Delbrück to characterize the onset of resistance in bacterial populations^22^. This assay exploits multiple replicates of clonal populations to bypass lineage-tracking issues and provides an elegant strategy to estimate mutation rates. In brief, parallel cell cultures are first expanded in non-selective conditions for a fixed number of generations, then exposed to a selective agent (originally a bacterial virus) and the number of cultures harboring growing mutants is counted. The fraction of resistant variants is then used to estimate the mutation rate through mathematical modeling taking into account multiple parameters, including proliferation rates and the total number of cellular divisions.

The fluctuation test has been previously modified to infer the acquisition of resistance to therapy in human tumors (e.g. by taking into account death rates)^23–26^, and adapted to describe evolutionary dynamics underlying development of resistance in clinically relevant conditions^27^. These versions of the fluctuation test are suitable to analyze the emergence and evolution of resistant cells already present in the tumor lesion before treatment initiation and to estimate the mutation rate of the cancer cell population in basal conditions. However, to the best of our knowledge, the available versions of the fluctuation test are not designed to quantify mutation rates in drug-tolerant persister cancer cells.

Here, we present a quantitative approach involving longitudinal biological experiments and mathematical modelling to characterize the transition of cancer cells to persistence under drug pressure and to measure mutability before and during therapeutic treatment. Specifically, we introduce a two-step fluctuation test to quantify mutations rates of CRC cells under standard growth conditions and during exposure to targeted therapy. The assay discriminates pre-existing resistant clones from *persister-derived* ones, allowing quantification of both spontaneous and drug-induced mutation rates.

## Results

### Growth dynamics of CRC cells in the absence of drug-treatment

With the final goal of calculating whether drug treatment affects the mutation rate of tumor cells, we first designed a set of biological experiments to infer growth dynamics parameters of CRC cell populations. These included: (i) growth rates in standard cell culture conditions (ii) population dynamics under treatment with targeted therapies and (iii) population dynamics of residual persister cells. To characterize the population of cancer persister cells that emerges in response to targeted therapies we studied two microsatellite-stable (MSS) CRC cell lines, for which we previously found evidence for the emergence of persisters^18^. DiFi cells carry amplification of the *EGFR* gene and are *RAS/RAF*-wildtype, which renders them highly sensitive to the anti-EGFR antibody cetuximab, paralleling the subset of CRC patients responsive to EGFR blockade^3,18^. As a second model we used *BRAF* V600E mutated WiDr cells, which are sensitive to BRAF inhibition in combination with cetuximab^18^, a therapeutic regimen recently introduced for the treatment of metastatic CRC patients with *BRAF* mutations^28,29^. These cell models were established before the clinical development of anti-EGFR and BRAF targeted therapies in CRC, ruling out the possibility that drug exposure could have occurred in the patients from which the cells were derived^30^.

To reduce the possibility that pre-existing resistant cells were present in the populations at the beginning of the assay, we first isolated individual clones for each cell model (namely WiDr cl. B7 and DiFi cl. B6), with growth and drug sensitivity profiles comparable to those of the parental population from which they originated (Extended Data Fig. 1a and b, respectively).

We first measured birth and death rates of clones in standard cell culture conditions. We defined WiDr and DiFi population dynamics by a standard birth-death process with two parameters: the birth rate *b*, i.e., the rate at which new cells are generated through cell division, and the death rate *d.* For each model (WiDr cl. B7 and DiFi cl. B6), we measured daily the total number of viable and dead cells (Extended Data Fig. 2a). The effective growth rate *(b – d)* was estimated with an exponential fit of the growth curve of each clone (Extended Data Fig. 2b and Extended Data Table 1); while the value *d/b* was estimated from the fraction of dead cells detected by flow cytometry (see Methods for mathematical details, Extended Data Fig. 2c and Extended Data Table 1). Finally, the values of the birth and death rate in the absence of treatment were obtained by combining the two estimates (values of *b-d* and *d/b*) (Extended Data Table 1).

### Growth dynamics of CRC cells under drug-treatment

The population dynamics of cancer cells during drug treatment involve both growth in presence of the drug and the transition of sensitive cells to a persister phenotype. The transition to persisters can be described as follows: each cell has the possibility to reversibly turn into persister at a certain rate, which could depend on the external conditions (e.g., drug concentration). Growth/division parameters of persister cells, as well as their death rate under drug treatment differ from those of the original sensitive population (see below for a mathematical formulation of these rules). To characterize the emergence of the persister subpopulation, and to quantify the intertwined processes of growth and transition to persister state upon drug treatment, we collected data from two sets of drug-response growth assays (Fig. 1a). In the first one (hereafter referred to as ‘*doses-response assay*’), we evaluated response of the two cell models to multiple doses of targeted therapies (Fig. 1a). Both DiFi and WiDr clones were exposed to increasing concentrations of the EGFR inhibitor cetuximab alone or in combination with the BRAF inhibitor dabrafenib, respectively. Cell viability of WiDr cells was assessed daily over a 5-day treatment period. Since we observed a slower decay in the DiFi cell number during cetuximab treatment, we extended measurements to 19 days in this model (Fig. 1a and 1b). Data from *doses-response assay* were used to analyze growth curves defined as number of live cells vs time and concentration of drug treatment (Fig. 1c, see Methods).

**Figure 1.**
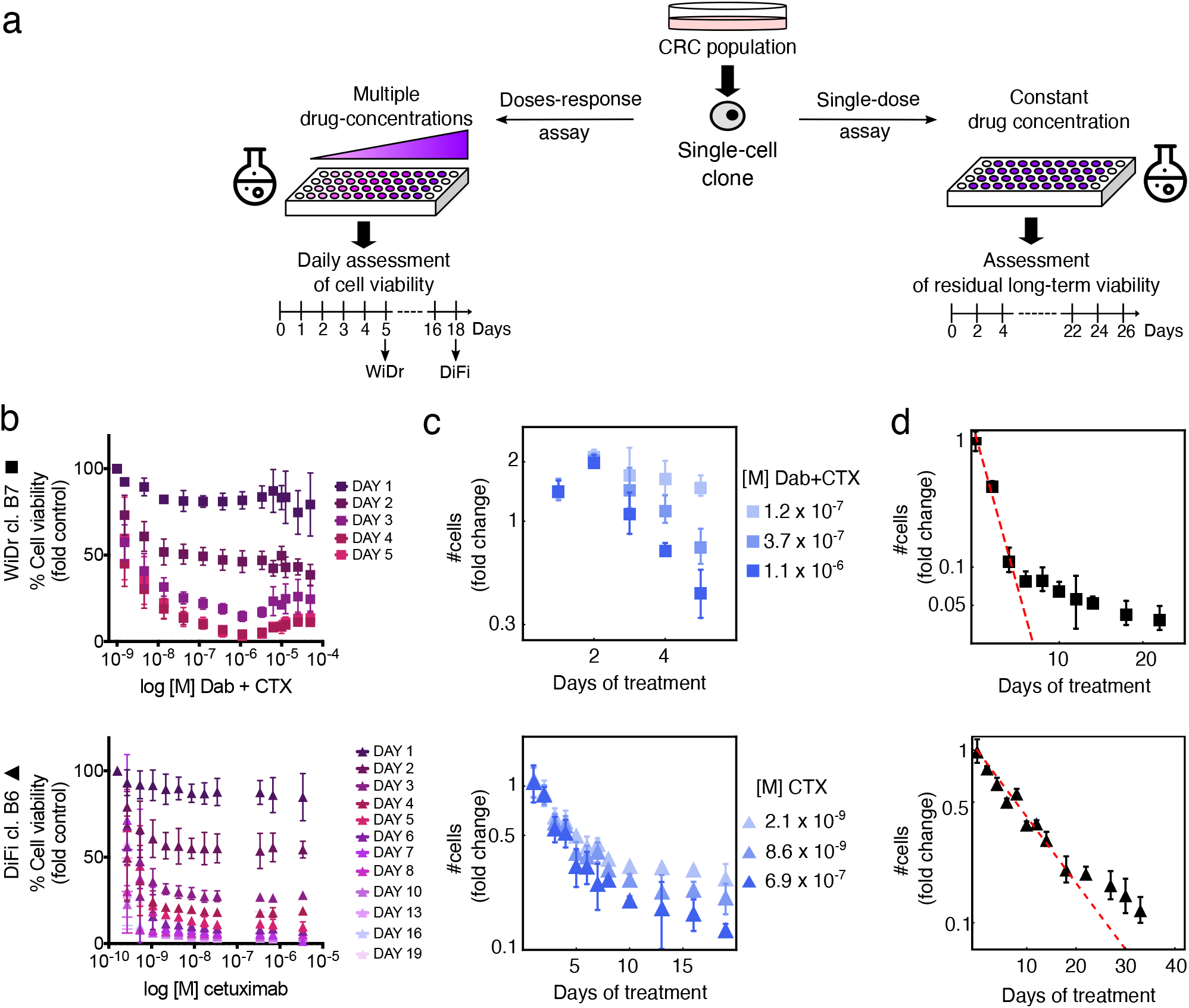
Population dynamics of CRC clones in response to targeted therapies. **a**, Schematic representation of the drug screening growth curve assays performed on CRC clones. **b**, In the *doses-response assay*, WiDr cells were treated with increasing concentrations of dabrafenib (Dab) + 50μg/ml cetuximab (CTX), while DiFi received increasing concentrations of cetuximab. Cell viability was measured by the ATP assay at the indicated time points. Results represent the average ± SD (n=3 for WiDr; n=5 for DiFi). **c**, Growth curves of the indicated cells under treatment, reported as foldchange of viable cells (log scale) vs time of drug exposure, were calculated from *doses-response assay* data, by normalizing cell viability at the indicated time points by the viability measured at day 0. Growth curves for three different drug concentrations for each clone are shown as average ± SD (n=3 for WiDr; n=5 for DiFi). **d**, Fold-change of viable cells (log scale, assessed by ATP assay) vs time of drug exposure for indicated cells in the *single-dose assay.* The total number of viable cells is compatible with an exponential decay with two-time scales, supporting the outgrowth of persisters (the dashed line indicates the initial slope). Symbols and error bars indicate means and standard deviations (n=2).

We then assessed response to a constant drug concentration by analyzing the fraction of surviving cells over 3 weeks of treatment (referred to as *‘single-dose assay*’) (Fig. 1a and d). The data highlight a biphasic two time-scale exponential decay (Fig. 1d), identified for bacteria as characteristic of the emergence of persister sub-populations^31^. We took this as a proof of the existence of persisters, and defined persisters as the residual sub-populations showing a very-slow death rate upon continuous treatment. Both DiFi and WiDr showed a slow but measurable decay in cell number, compatible with an exponential decay, suggesting a tendency of persisters to slowly die over time (Fig. 1d and Extended Data Fig. 3).

### The persister phenotype is induced by treatment of CRC cells with targeted therapies

To quantify the cell population dynamics during drug treatment, we developed a mathematical model of the transition of CRC cells to the persister state which we term *“transition to persisters”* or *TP* model. This model incorporates birth-death parameters and phenotypic switching in the deterministic limit (i.e., neglecting fluctuations due to stochastic demographic effects, see Methods)^32,33^. Fig. 2a summarizes the TP model dynamics. We exploited the model to quantify the transition rate and assess whether a sub-population of persister cells predated drug administration (and was therefore only selected by treatment pressure) or if the persister phenotype emerged at a measurable rate upon drug treatment. To this aim, the TP model considers three possible fates for drug-treated cells: i) death; ii) replication; iii) switching to persister state at a rate *λ* in the presence of the drug; the model further considers the possible pre-existence of an arbitrary steady fraction *f*_0_ of persisters (Fig. 2a).

**Figure 2.**
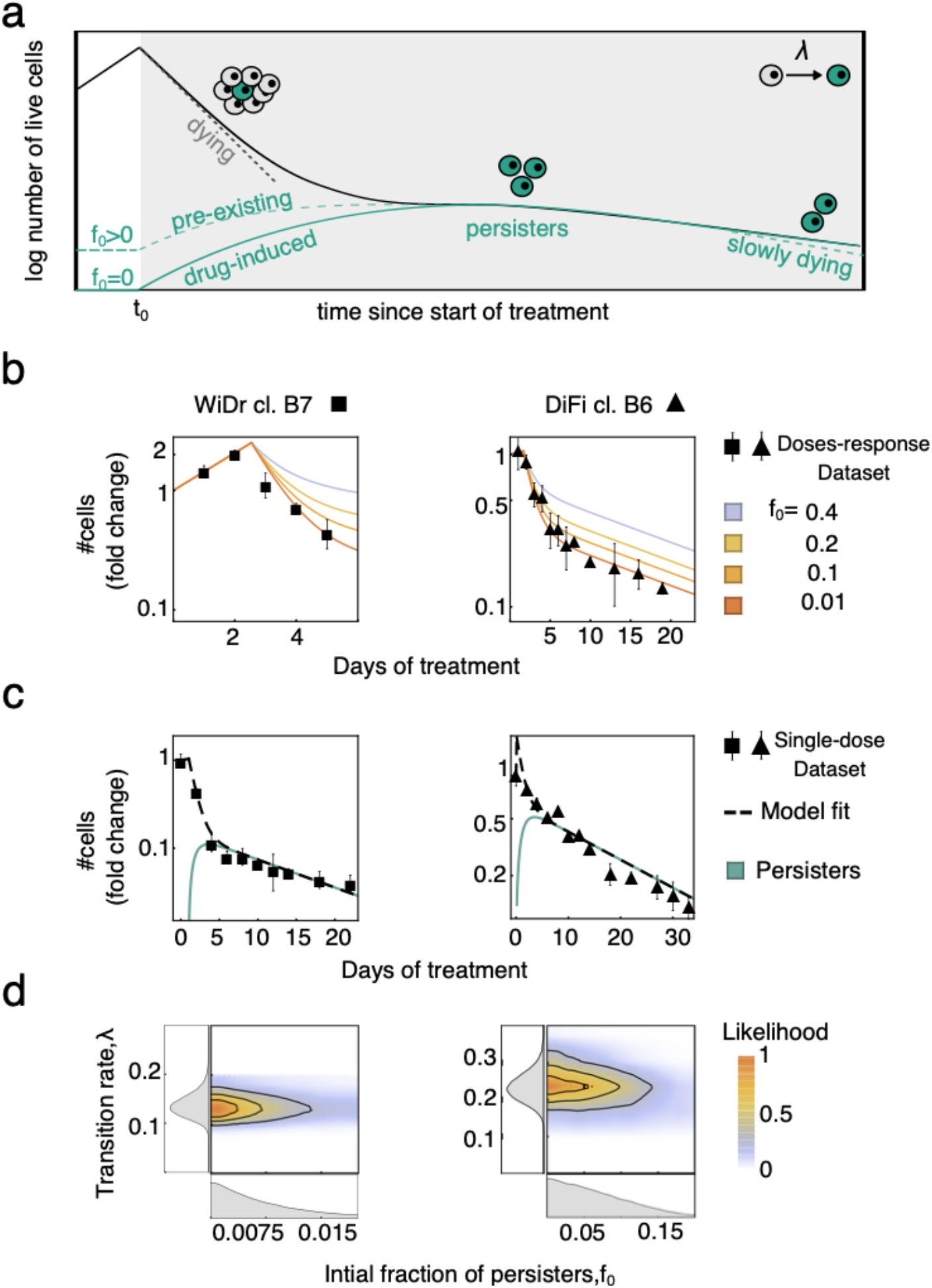
Targeted therapies induce the persister phenotype in CRC cells. **a**, Schematic representation of cell population dynamics under constant drug treatment. When cancer cells are exposed to targeted therapies, the number of viable cells starts to decline. A homogeneous population of sensitive cells (grey cells) would shrink exponentially to extinction (grey dashed line). Some cells survive drug treatment due to the transition to a persister phenotype (green cells, green lines) at a rate λ, and residual cells (solid black line) show a bi-phasic decay. A finite fraction of persister cells (*f*_0_) might be present in the population prior to drug treatment (*f*_0_ >0, dashed green line) or not (*f*_0_ =0, solid green line). Persisters cells show a reduced death rate during treatment, which results in a slow exponential decline of the cell population (green dashed line). **b**, Growth curves of CRC clones under treatment, calculated from the *doses-response assay*. Black symbols and bars represent averages ± SD of the *doses-response* dataset (n=3 for WiDr, n=5 for DiFi). Continuous lines indicate the TP model fit to the experimental data for different values of the initial fraction of persisters cells (*f*_0_, color coded). **c**, Fold-change of viable cells *vs* time of drug exposure for the indicated cells assessed based on the *single-dose* dataset. Black symbols and bars represent averages ± SD of the experimental data (n=2). The black dashed line indicates the fit of the TP model to the data, while the expected fraction of persisters cell is shown with the green solid line. **d**, Joint posterior distribution (contour plot, color coded with the normalized likelihood function) and marginalized posterior distributions (left and bottom panel, grey area indicates the Probability Density Function) of TP parameters describing the dynamics of persister cells: (i) the initial fraction of persisters cells (*f*_0_, bottom panel) and (ii) transition rate of sensitive to persister cells (*λ*) induced by the drug treatment. The likelihood function measures the agreement of the model to the experimental data as a function of the parameters considered.

The following equations define the dynamics of sensitive (X(t)) and persister cells (Z(t)) according to the TP model (under drug treatment):

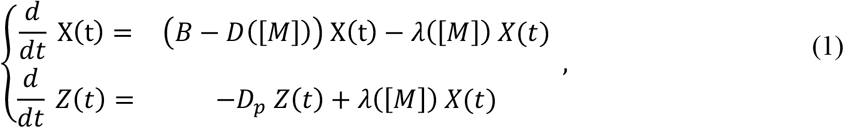

where *B* and *D*([*M*]) are, respectively, the birth and drug-dependent death rates of sensitive cells, while [M] is the drug concentration. Persister cells emerge with a drug-dependent transition rate *λ*([*M*]). Under drug treatment, persister cells die with rate *D_p_* > 0. The model assumes that persister cells that attempt to divide before developing drug-resistance mutations die (therefore a possible back-switching from persister to sensitive in presence of the drug would effectively contribute to the death rate). Further mathematical details of the TP model are given in the Methods.

The initial condition that specifies the solution to Eq. (1) is key for quantifying how much the transition to persister state is induced by the drug treatment. In particular, if the sensitive-to-persister transition is fully drug-induced, then untreated cells do not contain any persisters, i.e., 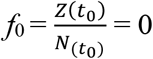.

Conversely, if some persisters pre-exist drug treatment, then the initial fraction of persister cells has a finite positive value (*f*_0_ > 0). If *f*_0_ is very small, some persisters may pre-exist the treatment, but the transition is mainly drug-induced. If instead *f*_0_ is comparable to the fraction of residual persisters after weeks of treatment, then the transition to persistence is not drug-induced.

To determine the most likely scenario, and to infer the TP model parameters, we used experimental data collected from drug-response growth curve assays (Fig. 1). Using results from the *doses-response assay* (focused on the population dynamics of the drug treatment at earlier timepoints), we defined parameters governing the dynamics of the model over a short timeframe, such as the initial fraction of persister cells *f*_0_ and the effective growth rate of treated cells.

Similarly, the *single-dose assay*, which follows the growth of CRC cells under treatment up to 30 days, was used to quantify model parameters that affect long-term dynamics, such as the transition rate of sensitive to persister cells *(λ)* and the effective death rate of persister cells (*D_p_*). By constraining model parameters from experimental data, we established which scenario would best describe the cell-based results.

Upon treatment, the number of cells started to decline within 1 to 3 days (*t*_0_), depending on the initial seeding density (Extended Data Fig. 4). The observed cell dynamics were coherent in experiments with different seeding densities once the growth curves were scaled (both in time and measured viability) to the maximum value reached at *t* = *t*_0_ (Extended Data Fig. 4).

The parameters of the TP model were then inferred with a standard Bayesian inference framework for both cell lines (Extended Data Table 2 and Methods for prior distribution used for the inference of the model parameters). DiFi displayed slower ‘dying’ dynamics compared to WiDr. In light of this, in WiDr we performed a joint fit of both the *doses-response* and *single-dose datasets*, while in DiFi we assessed growth curves in response to multiple doses of targeted therapies for up to 19 days, which allowed performing a model fit based on the *doses-response dataset* only (Methods).

We identified the best-fit TP model parameters given the experimental data, taking into account different values of initial number of persister cells (*f*_0_). We found that the best fit between the inferred TP model and experimental data occurs when *f*_0_=0, while the concordance decreases when *f*_0_ increases; we note that a *f*_0_ of 10% already significantly deviates from the data (Fig. 2b). Therefore, the TP model is consistent with the persister phenotype being predominantly drug-induced. In addition, the model properly describes the dynamics of *the single-dose assay* (Fig. 2c).

To further confirm the validity of the TP model, we next focused on the Bayesian statistics of the two model parameters describing the dynamics of persister cells: the transition rate λ and initial fraction of persisters *f*_0_. The joint posterior distribution of the Bayesian inference of these two parameters is shown in a heatmap that associates the likelihood of the TP model inference to the specific values of the two parameters (Fig. 2d and Extended Data Fig. 5). Importantly, we found that the transition rate to persistence λ estimated by the model fit does not vary when considering different values of the initial fraction of persisters *f*_0_ (Fig. 2d). The marginalized posterior probability of *f*_0_ is peaked at zero (Fig. 2d, bottom panel), and its upper bound is much smaller than the ratio between the persister population size (after all persister cells have emerged) and the total population size at the beginning of treatment. This implies that the inferred value of transition rate λ is independent from *f*_0_, and that the best concordance of the TP model to the experimental data is obtained for *f*_0_ =0.

Finally, to compare the scenarios *f*_0_=0 and *f*_0_>0, we used the Bayesian Information Criterion (BIC) and the Akaike Information Criterion (AIC), which are two standard Bayesian criteria used for model selection. According to both criteria, the TP model with *f*_0_=0 is preferred over *f*_0_ > 0 (Extended Data Fig. 6 and Extended Data Table 3, see Methods). We summarize the best-fit TP model parameters in Extended Data Table 4. These results support the concept that even if few persisters exist in the population before drug treatment, the majority of them must have transitioned to the persister phenotype after drug exposure.

### Cell abundance across wells is consistent with a drug-induced scenario for persister cells

We then performed an orthogonal experimental validation of the drug-induced scenario of persister state in both CRC cell models. Specifically, we measured how the number of persister cells varied across multiple wells, since the distribution of this parameter is expected to be different between a drug-induced and a pre-existent scenario^34^. To this aim, we seeded each cell model in multiple 96- well plates and quantified the distribution of persister cells (residual cell viability) among >400 independent wells after 3 weeks of drug treatment (Extended Data Fig. 7a). The observed abundance distribution across wells was consistent with a Poisson distribution (Extended Data Fig. 7b), supporting a drug-induced scenario in both CRC cell models, as confirmed by computer simulations (see^34^, where a similar method was used for mutational processes, and Extended Data Fig. 7c).

### CRC persister cells slowly replicate during drug treatment

Persisters are capable of surviving the lethal effect of the drug by entering a non-replicating or slowly replicating state^10^. We and others have recently reported that drug-treated cancer cells, alike bacteria exposed to antibiotics, initiate adaptive mutability stress response^18,20^ a process likely involving cell division and DNA replication. We therefore set out to establish whether persister cells divide under drug pressure. We treated both CRC cell models with the corresponding targeted therapies for two weeks, until cells reached the persister state, and then stained the residual persisters cells with Carboxy fluorescein succinimidyl ester (CFSE), a cell-permeable fluorescent dye allowing flowcytometric monitoring of cell divisions. Upon passive diffusion through the cell membrane of viable cells, CFSE is cleaved by intracellular esterases and retained within their cytoplasm. When a cell divides, CFSE is uniformly segregated between daughter cells; therefore, the reduction of CFSE intensity is proportional to cell divisions^35^.

CFSE-labelled DiFi and WiDr cells were analyzed by flow cytometry at different time points. To discriminate between persisters and pre-existing resistant cells, we seeded and treated each cell models in multiple 24-multiwell plates and excluded from the analysis wells containing clones that grew in two weeks in the presence of the drug. The analysis unveiled that in both CRC models, persister cells slowly replicate during treatment (Extended Data Fig. 8). These findings are in line with recent data showing a fraction of cycling persister cells emerging during treatment of lung cancer cells with an EGFR inhibitor^36^.

### Development of a fluctuation assay to quantify mutation rates during drug treatment

The measurement of mutation rates in the absence and in the presence of anticancer drugs required the development of a second mathematical model, which we refer to as “Mammalian Cells–Luria-Delbrück” or “MC-LD”. MC-LD is a fully stochastic birth-death branching process, describing the growth of resistant cells before and during drug treatment (Fig. 3a). We designate with *μ* the effective rate at which one individual (cell) becomes a mutant (therefore, resistant to treatment) and *μ_s_* and *μp* indicate mutation rates of sensitive (untreated) and persister cells, respectively (Fig. 3b, see Methods).

**Figure 3.**
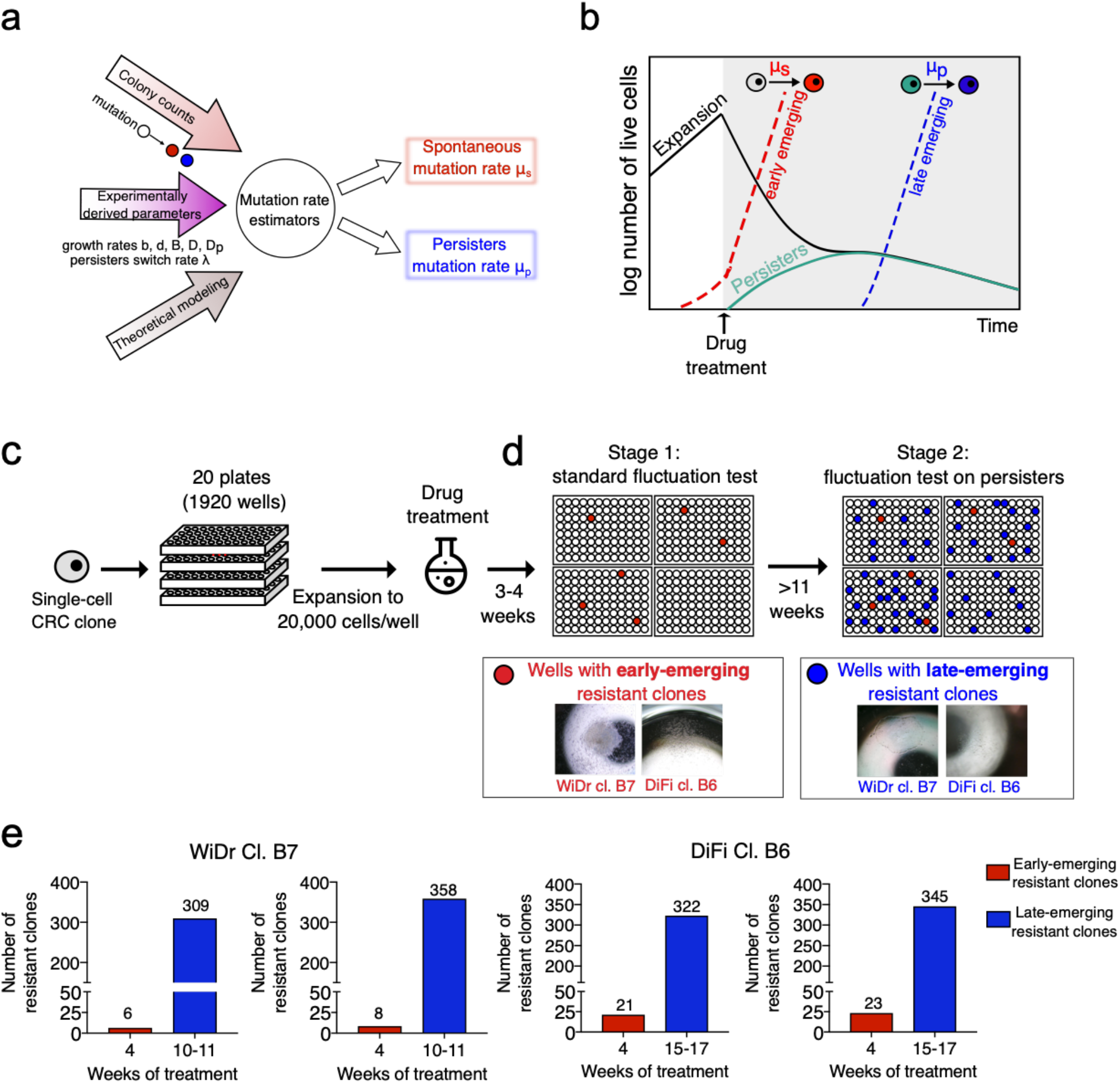
A modified Luria-Delbrück fluctuation test to measure mutation rates in mammalian cells. **a**, The modified fluctuation test, based on the inferred population dynamics and the MC-LD model, allows estimation of spontaneous (*μ_s_*) and persisters (*μ_p_*) mutation rates. **b**, Schematic representation of cell population dynamics of CRC cells during the fluctuation test. During the initial expansion in the absence of drug treatment CRC cells mutate with the spontaneous mutation rate (*μ_s_*). When cells are exposed to targeted therapies, pre-existing resistant cells are selected by the drug and give rise to early emerging resistant colonies (red dashed line), while sensitive cells start to decline in number (black solid line) and switch to the persister state (green solid line). Resistant cells derive from persister with a mutation rate *μ_p_* and give rise to late-emerging resistant colonies (blue dashed line). **c**, Schematic representation of the experimental assay underlying the fluctuation assay. WiDr and DiFi cells were seeded in twenty 96-multiwell plates, for a total of 1920 wells, and allowed to expand in the absence of drug for about 8 generations (reaching ~20000 cells/well). After the expansion, all the wells were treated with targeted therapy (100 μg/ml cetuximab for DiFi and 1μM dabrafenib + 50 μg/ml cetuximab for WiDr). **d**, Two sets of resistant clones were identified during the MC-LD experimental assay: the early-emerging resistant clones grown after 3-4 weeks (Stage 1), and the late-emerging resistant clones arising after >10 weeks (Stage 2) of constant drug treatment. **e**, Each bar graph lists the number of resistant clones counted at the indicated timepoints during MCLD experiment for each CRC clone. Red bars indicate early-emerging resistant clones (appearing in the first 4 weeks of drug treatment); blue bars indicate late-emerging resistant clones (appearing after ≥ 10 weeks of drug treatment). Results of two independent biological replicates for each clone are shown.

The MC-LD fluctuation assay is based on the experimental setting illustrated in Fig. 3c, d. WiDr and DiFi were seeded in twenty 96-multiwell plates and allowed to grow for a fixed number of cell divisions in standard culture conditions in the absence of drug (Fig. 3c); afterwards, a constant drug concentration was applied (Fig. 3d). The number of wells, the initial population size in each well and the time of cell replication in the absence of drug treatment were set by theoretical considerations using the MC-LD model, incorporating the population dynamics parameters we previously measured (Extended Data Tables 1 and 4).

In accordance with our previous work^8^, a small number of early-emerging resistant colonies were detected after 3-4 weeks of treatment (Fig. 3d, e; see Methods). In the vast majority of the wells, sensitive cells died, while drug-tolerant persister cells survived, as detected by measurement of residual cell viability (Extended Data Fig. 7a)^8^. Constant treatment of the residual persister cells was then continued to perform the second step of the fluctuation assay (Fig. 3d); after several weeks of treatment, late-emerging resistant colonies appeared in a subset of the wells in which persister had previously been detected (Fig. 3d, e).

We ran multiple MC-LD model simulations, with input parameters inferred with the TP model, and found that resistant clones emerging at late time points (>4 weeks of treatment), are extremely unlikely to originate from pre-existing resistant cells (Fig. 4a and Extended Data Fig. 9). In accordance with previous work^8^ we considered the resistant colonies that became microscopically visible within the first 4 weeks of drug treatment (early-emerging resistant) as those representing the pre-existing resistant cells, that is the mutant cells that emerged during the expansion phase by spontaneous mutation. We reasoned that resistant colonies that slowly emerged after ≥10 weeks of drug treatment (late-emerging resistant) in persisters-containing wells could have developed drug resistance mutations during the adaptive mutability process which we and other have shown to occur in persister cells^18,20^ (Fig. 4a and Extended Data Fig. 9).

**Figure 4.**
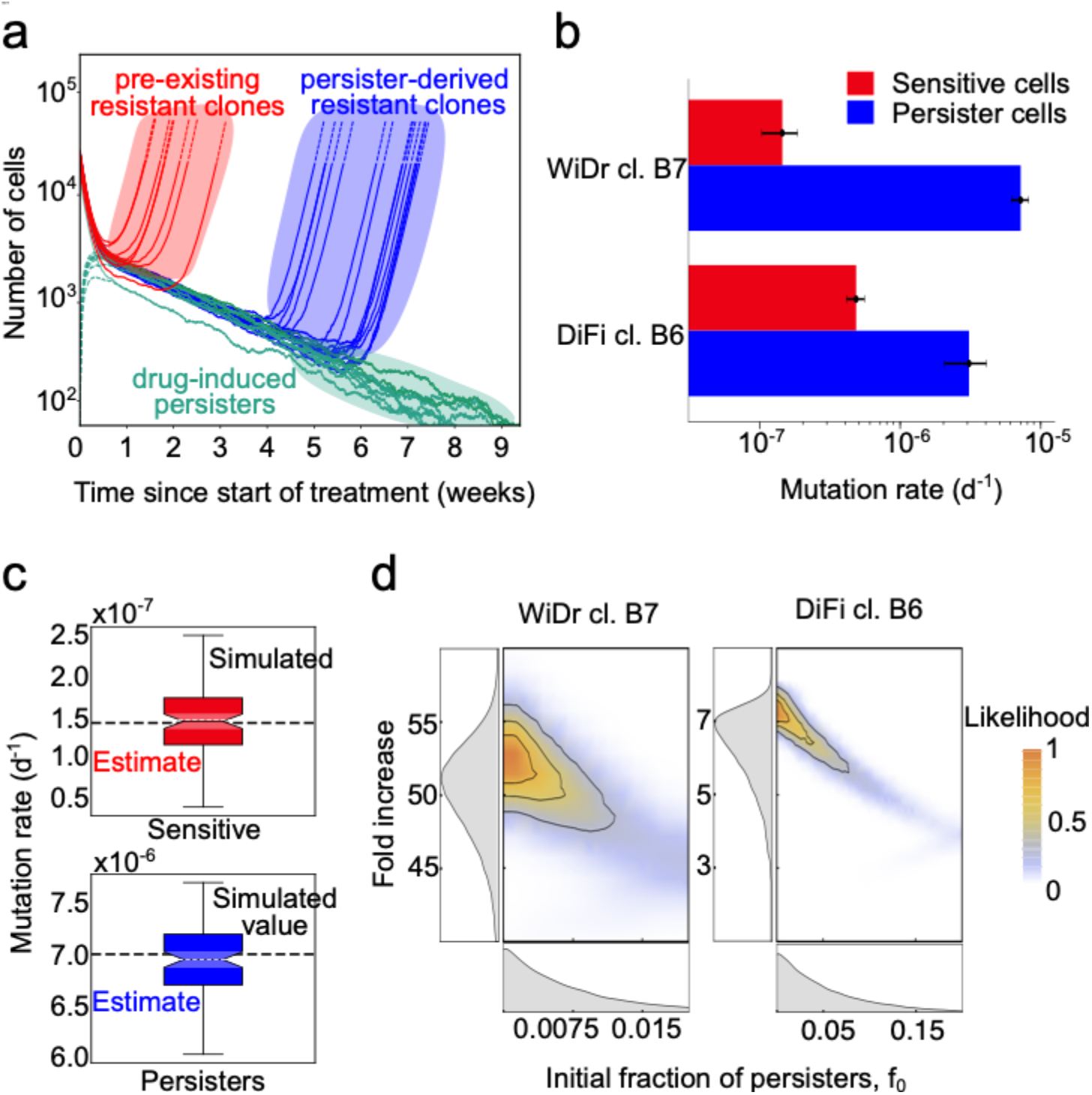
Quantification of mutation rates in persister cells. **a**, Simulated data for the assay described in Fig. 3. The experimentally measured MC-LD model parameters and the model-derived estimators of mutation rate, for sensitive and persister cells, were used to simulate the time of appearance of pre-existing and persisters-derived resistant cells. **b**, Quantification of mutation rates for sensitive (red) and persister (blue) cells in the MC-LD experiment. The indicated cell models were seeded and treated as described in Fig. 3. Mutation rates were calculated from the experimental data based on population parameters and the number of pre-existing (early-emerging) and persisters-derived (late-emerging) resistant clones as described in Fig. 3. Results represent inferred mutation rates (for sensitive and persisters cells for each clone) with bar plots showing mean and the 99% C.I. of the posterior distributions of the mutation rates (n=2). **c**, Validation of mutation rates estimator with model simulations. The box plots represent the distribution of the estimated mutation rates for 100 runs of the entire experiment using the parameters reported in Extended Data Table 1 and 4. The dashed lines represent the input value of the mutation rate used in the simulation. **d**, Joint posterior distribution (contour plot, color coded with the normalized likelihood function) and marginalized posterior distributions (left and bottom panel, grey area shows the Probability Density Function) of (i) the initial fraction of persisters cells (*f*_0_, bottom panel) and (ii) fold increase of the mutation rate of persister cells compared to mutation rate of sensitive cells (*μ_p_*/*μ_s_*). The likelihood function measures the agreement of the model to the experimental data as a function of the value of the parameters considered.

As in a standard fluctuation test, the mutation rate can be inferred from the observed fraction of wells containing resistant cells. In the model, this fraction corresponds to the expected probability of observing a resistant clone in a well in a given time interval [0,*T*]. To compute this probability in the MC-LD model, we assumed that resistant cells divide with rate *b* and die with rate *d*, just like untreated cells. Because of reproductive fluctuations (genetic drift), cells carrying drug-resistance mutations can still go extinct, and only a fraction of the mutants, which we refer to as “established mutants”, survive stochastic drift. The probability of surviving stochastic drift in a time interval Δ*t*, denoted here as *ψ*(Δ*t*), is a well-known result of the birth-death process^37,38^ (see Methods).

We derived analytically an approximate solution of the model, by considering that the number of mutant cells established in the time interval [0,*T*] follows a Poisson distribution with expected value 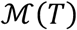. Consequently, the probability of having at least one mutant is given by

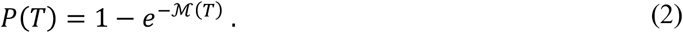

In order to quantify the spontaneous mutation rate of cancer cells (i.e., before drug administration), we focused on the resistant cells established by the time *T_treat_* before treatment administration. The expected number of resistant cells that emerged from sensitive cells in this time interval reads:

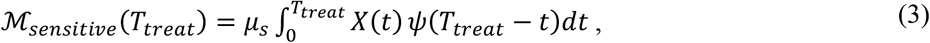

where *μ_s_* is the the mutation rates of sensitive cells.

To quantify the mutation rate of persister cells *μ_p_*, we consider resistant cells that emerged by the time *T* since the beginning of the drug treatment. The expected number of resistant cells emerged from persister cells reads:

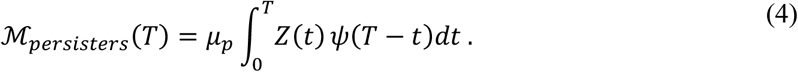

We emphasize that Eq. (2–4) are connected to the solution of the TP model, Eq. (1). Hence, the solution of the MC-LD model is defined in terms of the same parameters that were estimated above with the TP model (see the Methods for further details).

We then used this solution of the MC-LD model to derive estimators of mutation rates of sensitive cells *μ_s_* (encompassing the fraction of wells with early-emerging resistant cells) and of persisters cells *μ_p_* (corresponding to the fraction of wells with late-emerging resistant clones).

### Persister CRC cells show increased mutation rate under drug treatment

Data collected with the two-steps MC-LD fluctuation test allowed inferring mutation rates of sensitive (*μ_s_*) and persisters (*μ_p_*). Since accurate quantification of slow cell division in persister cells was unfeasible, we evaluated the mutational processes as chronological (measured in mutations per day) rather than replicative (mutations per generation). This choice is conservative, as untreated cells divide in any case much faster than persisters, hence the ratio between replicative mutation rates of cells displaying the two phenotypes must be in any instance higher than for chronological rates.

Notably, we found that mutation rates were increased by a factor of 7- to 50-fold in cells that survived and tolerated for several weeks doses of targeted therapies that were instead lethal for the majority of the parental population (Fig. 4b and Extended Data Table 5). This result was consistent across multiple biological replicates, both in DiFi and WiDr cells and in response to either EGFR blockade or EGFR/BRAF concomitant inhibition, respectively (Fig. 4b and Extended Data Table 5). To further validate the consistency of the mutation rate inference based on the MC-LD model, we ran multiple simulated replicates of the experiment, using a set of sensitive (*μ_s_*) and persister (*μ_p_*) mutation rates, and we then used the MC-LD estimators on the synthetic data. Fig. 4c compares boxplots of the estimated mutation rates across replicates of simulated experiments with the actual values of mutation rates used as inputs to the simulations. The agreement between these values validates our estimates.

We further reasoned that the value of the initial fraction of pre-existing persister cells *f*_0_ could play a role in the estimate of the mutation rate. We therefore assessed whether and to what extent the inferred value of the mutation rate is affected by the presence of different amounts of pre-existing persister cells using our estimators within a Bayesian framework (Fig. 2d). This approach returns the mutation rate, considering a range of realistic values of *f*_0_, and their probability. As a result of this analysis, we obtained the fold increase of the mutation rate of persister cells as a function of *f*_0_, in the entire range of values that are compatible with the dynamics observed in the growth curve assays experimentally assessed in Figs. 1 and 2. Based on this, Fig. 4d summarizes the results of this inference in a joint heatmap of *f*_0_ and the fold increase of persisters mutation rate compared to that of sensitive cells. We found that considering all representative values of *f*_0_ that are compatible with our experimental data the increase of mutation rate in persister cells remains strongly supported.

To summarize these findings, we propose a quantitative model for the evolutionary dynamics of CRC cells exposed to clinically relevant concentrations of targeted therapies (Fig. 5). In non-stressful conditions, cancer cells replicate and spontaneously acquire mutations which can confer resistance to targeted therapies (pre-existing resistant mutations) at a replicative mutation rate *μ_s_*. However, when cancer cells are exposed to the hostile environment generated by targeted therapies, the majority of sensitive cells quickly die while a subset of parental cells switch to a long-lasting surviving persister state at a rate λ and in a drug-induced manner. Previous and current findings indicate that persister cells, under constant exposure to lethal doses of drugs, initiate a stress response which affects DNA replication fidelity^18,20^, thus leading to a measurable increase of their mutation rate (*μ_p_*), therefore rising the probability that alterations conferring drug resistance could occur.

**Figure 5.**
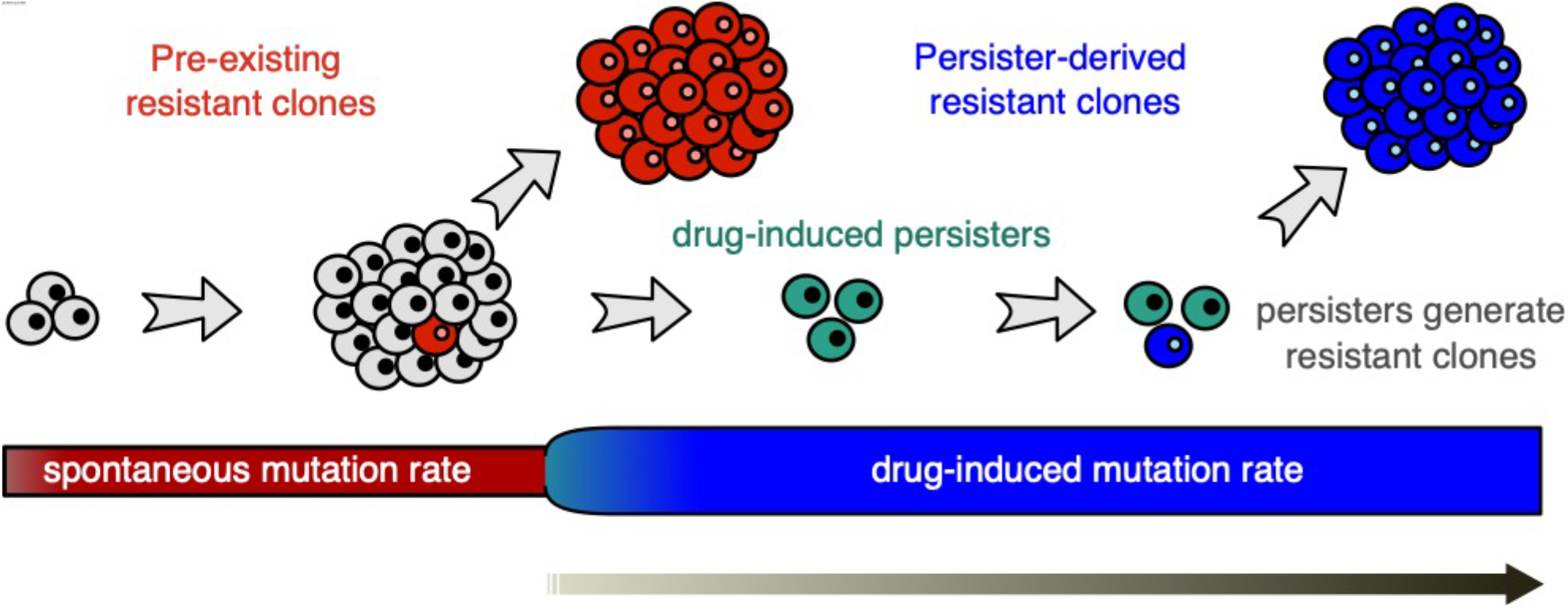
Schematic representation of CRC cells mutational dynamics during drug treatment. Untreated cells spontaneously acquire resistant mutations at a replicative spontaneous mutation rate *μ_s_*. When cancer cells exposed to targeted agents, a surviving persister phenotype is induced in a drug-dependent manner. Persister cells under constant drug exposure reduce DNA replication fidelity and increase their mutation rate at rate *μ_p_*. This in turn boosts genetic diversity and favors the emergence of resistant clones driving tumor recurrence and treatment failure.

## Discussion

A prevalent view is that acquired resistance to inhibitors of oncogenic signaling is driven by mutant (drug resistant) cells that are already present in the tumor mass before treatment initiation^3,4^ This concept has significant impact on clinical treatment of cancer patients, as it implies that resistance is a ‘fait accompli’; the time to recurrence is therefore simply the interval required for pre-existing drug resistant cells to repopulate the lesion. To limit the impact of drug resistant cells, novel drugs (such as the EGFR inhibitor osimertinib in NSCLC) were introduced and combinatorial regimens are being considered over monotherapy approaches with the goal of eradicating pre-existing resistant cells or reducing the probability that they will lead to treatment failure^27,39,40^.

Interestingly, in a considerable subset of patients targeted therapies lead to long lasting reduction of tumor burden, and relapse only occurs after prolonged disease stabilization. Such a situation challenges the current view, and several pieces of evidence support a role of persister cells in treatment relapse^12^. For example, we and others have recently shown that adaptive mutability fosters acquisition of mutations driving resistance by increasing genomic instability^18,20^. However, the lack of models to quantitatively characterize the behavior of persisters under treatment has so far hampered progress in the field, including the possibility to actually measure mutation rates of tumor cells during treatment. Our approach integrates experimental data and mathematical modelling to investigate population dynamics of tumor cells exposed to targeted therapies. We developed a two-step fluctuation assay, which allows quantitative comparisons of spontaneous (basal) and drug-induced mutation rates in cancer cells.

In agreement with previous observations, we found that when a population of cancer cells is constantly exposed to targeted therapies, a subset of drug-tolerant persister cells emerges^7,8,13,15^. While previous observations of cancer persister cells were mostly qualitative, we describe a quantitative definition of cancer persisters based on the characterization of long-term response of CRC cells to targeted therapies^19,31^. Persisters were firstly and extensively described in bacteria in response to antibiotic treatment, where the hallmark of antibiotic persistence is the occurrence of biphasic killing curves^31^. Indeed, our analysis of CRC cells dynamics during long-term drug exposure points to a biphasic killing curve, characterized by a rapid decline of sensitive cells and a slow transition to persistence (Fig. 1d). Such a biphasic killing curve, building on the previous evidence^18^ that these cells generate, upon drug withdrawal, a population that is again sensitive to the same targeted therapy (and develop new persisters after a second exposure), can be taken as the proof of the emergence of a persister sub-population in CRC cells upon EGFR and BRAF inhibition.

Mathematical modeling of population dynamics under drug exposure indicates that the persister phenotype is largely drug-induced in CRC cells. Although we cannot exclude that in a tumor lesion a few persisters might already exist, the TP model, together with computational and experimental validations, strongly supports the notion that drug-induced sensitive-to-persister transition is a prevalent path to the development of this phenotype. This conclusion is in line with recent evidence of a chemotherapy-induced persister state described in CRC^15^. Whether all cancer cells have equal capacity to transition to persister or if a fraction of them is predetermined/predisposed to that, as suggested by a recent unpublished report^36^, remains to be established.

Importantly, we show that even if we consider the presence of a small subset of pre-existing persister cells, this possibility does not affect our conclusion that mutation rates increase under treatment. In essence, our mathematical models and the experimental evidence indicate that persisters slowly replicate during drug exposure and significantly increase their mutation rate, independently from the initial fraction of persister cells.

Based on our experimental data and models (Fig. 4a and Extended Data Fig. 9), we assumed that late-emerging resistant clones derive mostly from persister cells. If all treated cells increased their mutation rate under drug treatment, the contribution of sensitive cells to resistance would be exhausted after few weeks of treatment, as we show that they go extinct within a few days (Fig. 1d). Instead, new resistant clones keep emerging after several (10-20) weeks of continuous drug treatment (Fig. 3).

The evidence of active cell cycle progression alongside to increase of mutagenic rate in persister cells further supports our previous findings of ongoing adaptive mutagenesis under drug-induced stress fostering acquisition of resistance^18^. The Mammalian Cells–Luria-Delbrück (MC-LD) approach used here to quantify the effect of targeted therapies on the mutation rate of cancer cells could in principle be applied to measure whether and how a wide range of environmental conditions affect persister phenotype and mutation rates in mammalian cells.

For example, it would be interesting to deploy the same strategy to systematically identify chemical or physical agents affecting mutations rates in human cells. Note that the treatments used here do not induce DNA damage directly, though they could indirectly, as we previously showed^18^. Importantly, our analysis is not only limited to mutation rates, but also allows assessing whether drugs or environmental conditions restrict or reverse the transition to the persister state, rendering cells sensitive again to treatment.

The combined experimental and modeling framework presented here may have broad and far-reaching clinical implications. The emergence of persister cancer cells and the increase of mutation rate during adaptive mutability stress-response^18^ could be particularly relevant when targeted therapies lead to an initial extensive reduction of tumors mass, followed by long disease stabilization and eventually to relapse^5,6^. Furthermore, it would be of interest to assess whether what we observed in CRC cells also occurs in other tumor types in which agents targeting EGFR and BRAF are commonly used such as melanoma and lung cancer^41^. Equally important will be assessing whether exposure of cancer cells to agents directed against other oncogenic targets, such as HER2, NTRK or ALK, leads to similar phenotypic and mutational routes.

In conclusion, our data suggest that commonly used anticancer therapies trigger persistence and increase mutation rates in tumor cells and that innovative therapeutic strategies could be exploited to impair the emergence of persistence and resistant mutations, potentially extending the efficacy of clinical treatments.

## METHODS

### Experimental setup and data collection

#### Cell cultures

Cells were routinely supplemented with FBS 10% 2mM L-glutamine, antibiotics (100U/mL penicillin and 100 mg/mL streptomycin) and grown in a 37°C and 5% CO_2_ air incubator. Cells were routinely screened for absence of Mycoplasma contamination using the Venor® GeM Classic kit (Minerva biolabs). The identity of each cell line was checked no more than three months before performing the experiments using the PowerPlex® 16 HS System (Promega), through Short Tandem Repeats (STR) tests at 16 different loci (D5S818, D13S317, D7S820, D16S539, D21S11, vWA, TH01, TPOX, CSF1PO, D18S51, D3S1358, D8S1179, FGA, Penta D, Penta E, and amelogenin). Amplicons from multiplex PCRs were separated by capillary electrophoresis (3730 DNA Analyzer, Applied Biosystems) and analyzed using GeneMapper v.3.7 software (Life Technologies).

#### Isolation of CRC-derived clones

CRC clones were obtained by seeding WiDr and DiFi CRC populations at limiting dilution of 1 cell/well in 96-multiwell plates in complete medium. Clones were then selected for having growth kinetics and drug sensitivity comparable to the parental counterparts. For growth testing, WiDr and DiFi populations and derived clones were seeded in 96- multiwell plates (2×10^3^ cells/well and 3×10^3^ for WiDr and DiFi respectively) in complete medium. Plates were incubated at 37°C in 5% CO_2_. Cell viability, assessed every day for 4 days by measuring ATP content through Cell Titer-Glo® Luminescent Cell Viability assay (Promega), was compared to cell viability assayed at day 1. For drug sensitivity testing, cells were seeded at different densities (2×10^3^ cells/well and 3×10^3^ for WiDr and DiFi respectively) in medium containing 10% FBS in 96- multiwell plates at day 0. The following day, serial dilutions in of the indicated drugs were added to the cells in serum-free medium (ratio 1:1) in technical triplicates, while DMSO-only treated cells were included as controls. Plates were incubated at 37°C in 5% CO_2_ for the indicated time. Cell viability was assessed by measuring ATP content through Cell Titer-Glo® Luminescent Cell Viability assay (Promega). Dabrafenib was obtained from Selleckchem. Cetuximab was kindly provided by MERCK.

#### Growth rates of CRC cell clones before drug treatment

Clonal spontaneous growth is defined by the following parameters: the rate at which cells are born (birth rate, *b*), the rate at which cells die (death rate, *d)* and the net growth rate *b-d.* To estimate the *b-d* rate, CRC cell clones were seeded at 3.5-4×10^4^ cells/well in 6-multiwell plates. Plates were incubated at 37°C in 5% CO_2_. Starting from the following day, the number of viable cells was assessed by manual count in trypan blue 0.4% (Gibco^TM^) by two operators independently at the indicated time points, in order to obtain the clones’ net growth rate. To estimate *d/b*, cells were seeded at different densities (3.5-4×10^5^ cells/well) in multiple 6-multiwell plates. Plates were incubated at 37°C in 5% CO_2_. At each time point, cells were collected and stained with Propidium Iodide (Sigma Aldrich) following manufacturer’s instructions. Cells were then analyzed by flow cytometry. The cells in sub-G1 phase were considered dead and used to estimate *d/b.* The values of birth (*b*) and death rate *(d)* were then obtained by combining *b-d* and *d/b* estimates (see below).

#### Doses-response growth curve assay

CRC cell clones were seeded at different densities (2×10^3^ cells/well and 3×10^3^ for WiDr and DiFi respectively) in medium containing 10% FBS in multiple 96- multiwell plates at day 0. The following day, serial dilutions of the drugs were added to the cells in serum-free medium (ratio 1:1) in technical triplicates, while DMSO-only treated cells were included as controls. Cell viability of WiDr and DiFi clones was assessed at indicated time points over 5 and 18 days of constant treatment, respectively, by measuring ATP content through Cell Titer-Glo® Luminescent Cell Viability assay (Promega).

#### Single-dose growth curve assay

DiFi and WiDr CRC cell clones were seeded in multiple 96- multiwell plates at 1000 or 500 cells/well, respectively. Cells were allowed to expand for a fixed number of generations until a population size of 10000-20000 cells/well was reached. At that point, treatment was added (100 μg/ml cetuximab for DiFi and 1μM dabrafenib + 50μg/ml cetuximab for WiDr). Plates were then incubated at 37°C in 5% CO_2_ and cell viability was assessed at the indicated time points by measuring ATP content through Cell Titer-Glo® Luminescent Cell Viability assay (Promega) over 22 and 32 days of constant treatment (for WiDr and DiFi respectively). Medium and treatment were renewed once a week. To test the effect of different seeding densities on the residual viability assayed, each clone was seeded at different densities (3-20×10^3^ cells/well) in complete medium. The following day, treatment was added (100 μg/ml cetuximab for DiFi and 1μM dabrafenib + 50μg/ml cetuximab for WiDr) and viability was assessed at the indicated time points by measuring ATP content.

#### Staining with Carboxy fluorescein succinimidyl ester (CFSE)

CRC clones were seeded at 2.5×10^5^ (WiDr) and 6.5×10^5^ (DiFi) cells in multiple 10cm dishes. The following day, untreated cells were stained with CellTrace™ CFSE Cell Proliferation Kit (Invitrogen™) according to manufacturer’s instructions. At the indicated timepoints, starting from the day after staining (T0), cells were collected and resuspended in 1mL PBS with Zombie Violet™ 1000x (BioLegend®) to exclude dead cells. Cells were then analyzed by flow cytometry. For persisters proliferation analysis, CRC clones were seeded at 2×10^4^ cells/well in several 24-multiwell plates. The following day, cells were treated with 100 μg/ml cetuximab (for DiFi) or 1μM dabrafenib + 50μg/ml cetuximab (for WiDr) and incubated at 37°C in 5% CO_2_ for 14 days (renewing treatment after 1 week) until a population of persister cells emerged in each well. Then, cells were stained with CellTrace™ CFSE Cell Proliferation Kit (Invitrogen™) according to manufacturer instructions. At the indicated timepoints, starting from the day after staining (T0) plates were checked to exclude from the analysis wells containing resistant clones. Cells from the remaining wells were collected and resuspended in 1mL PBS with Zombie Violet™ 1000x (BioLegend®) to exclude dead cells and analyzed by flow cytometry. Medium and treatment were renewed once a week during all the time of experiment. Flow cytometry was performed using the FACS Dako instrument and analyzed with a Python script based on standard libraries (FlowCal, FlowKit). The following gating strategy was used. First, cells were selected with a light scattering gate (FSLin vs SS), excluding cell doublets with a single cell gate (FSArea vs SSArea). The following cutoffs were used: (i) *FSLin:* lower 5000 and upper 60000; (ii) *SSLin:* lower 3000 and upper 63000; (iii) *FS Area:* lower 3000 and upper 60000; *SS Area:* lower 2000 and upper 63000. We then evaluated the bi-dimensional distribution of the remaining data points in the space of the coordinates FS Area and SS Area, and retained all the data-points that were included in the 99nth percentile of the distribution. Viable cells were selected by excluding Zombie Violet™-positive cells and CFSE signal was detected by measuring Fitc signal.

#### Characterization of distribution of persister cells

DiFi and WiDr cell clones were seeded in multiple 96-multiwell plates at 1000 or 500 cells/well, respectively. Subsequently, cells were allowed to expand until they reached 10000-20000 cells/well. Cell viability was then assessed by measuring ATP content to normalize for cell number prior to treatment initiation. The remaining plates were treated with targeted therapies (100 μg/ml cetuximab for DiFi and 1μM dabrafenib + 50 μg/ml cetuximab for WiDr). Medium and treatment were renewed once a week. After 3 weeks of constant drug treatment, residual viability was assessed by measuring ATP content through Cell Titer-Glo® Luminescent Cell Viability assay (Promega).

#### Two-steps fluctuation assay

DiFi and WiDr clones were seeded at 1000 or 500 cells/well, respectively, in twenty 96-multiwell plates each, for a total of 1920 independent replicates. Cells were allowed to expand for a fixed number of generations until they reached 10000-20000 cells/well. Next, treatment was administered (100 μg/ml cetuximab for DiFi and 1μM dabrafenib + 50 μg/ml cetuximab for WiDr). Plates were incubated at 37°C in 5% CO_2_ for the indicated time. Media and drug treatment were renewed once a week. After 3-4 weeks of treatment, pre-existing resistant colonies were clearly distinguishable at the microscope and counted by two independent observers. The number of pre-existing resistant clones was used to estimate the spontaneous mutation rate of CRC clones (see methods section *Estimator of mutation rate for sensitive cells* below). After 10-11 weeks, resistant colonies started to emerge in wells where only persisters were previously present. The number of persister-derived resistant clones was used to estimate the mutation rate of persister cells under constant treatment (see methods section *Estimator of mutation rate for persister cells* below). Pictures of the resistant colonies were acquired using a ZEISS Axio Vert. A1 microscope equipped with a True Chrome HD II camera.

### Theoretical Modeling

#### Deterministic and stochastic model

In this study we have developed and used two distinct mathematical models to investigate the dynamics of cell populations. The first model describes the transition to persister state (“TP model”), and is a birth-death model with phenotypic switching, which we explored in the deterministic limit, i.e., neglecting statistical fluctuations and only considering expected values of the model output. We made use of this model for two purposes: (i) to infer the parameters of persisters dynamics, and (ii) for the Bayesian model-selection inference procedure that was used to infer whether the transition to persisters is drug-induced or not. This deterministic model is described in the sections *Dynamics of sensitive (untreated) cells* and *Dynamics of cell population under drug treatment* below.

The second model, which we named *Mammalian Cells-Luria Delbrück* or “MC-LD” model, is a fully stochastic birth-death branching process that includes the mutational processes of sensitive (untreated) and persister cells (under treatment). In order to measure the mutation rate, stochastic fluctuations cannot be neglected. Therefore, we considered this extension of the TP model including (i) stochastic fluctuations around the expected values of the deterministic limit and (ii) the mutational processes of both sensitive and persister cells. Relatedly, we did not include the latter process in the TP model, since during first days of drug treatment death of sensitive cells or transition to persisters are predominant, while acquisition of mutations driving resistance becomes significant only after weeks of treatment.

We have derived an approximate solution for the fraction of resistant wells in the MC-LD model, to derive estimators of mutation rate, as described in the methods section *Inference of the mutation rate from a two step fluctuation assay* below, and we have run the model by direct simulation in order to design the biological experiments and validate the estimators that we obtained (Fig. 4a, c). Simulations of this model were also used to investigate the distribution of the number of persisters cells after 3 weeks of treatment (Extended Data Fig. 7), by setting to zero the value of the mutation rates.

#### Dynamics of sensitive (untreated) cells

The drug sensitive clonal cell populations were modelled with a standard birth-death process. Their dynamics is therefore described by two parameters: (i) the birth rate (*b*) (i.e., the rate at which new cells are generated by replication) and (ii) the death rate *(d).* This model describes an exponential growth for the number *N_s_*(*t*) of viable cells at time *t*:

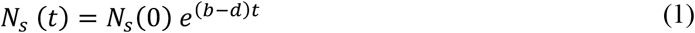

where *N_s_* (0) is the number of viable cells present at *t* = 0. Another dynamical variable which can be measured from the flow cytometry analysis (Extended Data Fig. 2c) is the fraction of dead cells *δ* which, at equilibrium, takes the value

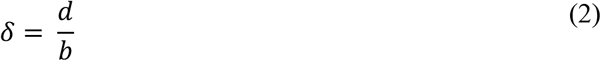

Hence, an indirect estimate of *b* and *d* can be obtained in two steps: (i) an exponential fit of the observed growth curves (Eq. 1) gives the value *b* - *d*; (ii) an estimate of the asymptotic fraction of dead cells (Eq. 2) gives the value *d/b.* We do not model explicitly the possibility of reversible switching to persister state in absence of treatment, as this process is not observable in our experimental data. However, the model accounts for this process via the parameter *f*_0_, the fraction of persister cells at the time of treatment (see below).

#### Dynamics of cell population under drug treatment

We focus on the dynamics of a population of *N*(*t*) = *X*(*t*)+*Z*(*t*) total viable cells, consisting of a combination of *X*(*t*) sensitive cells and *Z*(*t*) persister cells. The population is assumed to grow in presence of a drug with constant drug concentration [*M*]. Treated sensitive cells can (i) reproduce with a birth rate *B*, (ii) die with a drug-dependent death rate *D*([*M*]), (iii) switch to the persister state with a drug-dependent transition rate *λ*([*M*]).

Persisters cells display a moderate division rate under drug treatment (b≃0.3 days^-1^, Extended Data Fig. 8), and their observed dynamics under long-term treatment indicates a slow decline in cells number (Fig. 1d), compatible with a negative, but small, effective growth rate. These two observations can be described by a scenario where persister cells that attempt to divide before developing drug-resistance mutations die out (i.e., for which the net growth rate is ≃ 0), while nondividing persister cells slowly die during the treatment. In this scanario back-switching from persister to sensitive in presence of the drug effectively enters the model as a contribution to death rate, and this combined dynamics are described by the effective death rate *D_p_* > 0.

Drug effect is assumed to be delayed by a time *t*_0_ after the administration of the drug to the cancer cell population. During this time interval, the model assumes that the cell popoulation grows with a net growth rate *S*_0_:

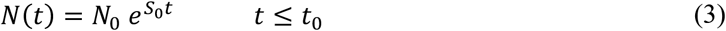

Where *N*_0_ is the initial number of cells. The quantity 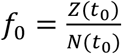 is the fraction of persister cells present at the time *t*_0_, i.e., at the time of the effective initiation of the drug effect. This parameter effectively incorporates the possibility of pre-existence of persisters due to reversible switching in absence of the drug. Under these assumptions, the dynamics of the fraction of sensitive cells during treatment is described by

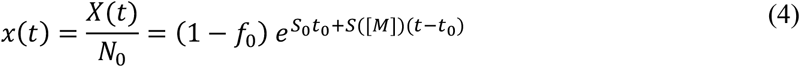

and we have defined *S*([*M*]) = *B* — *D*([*M*]) — *λ*([*M*]) as the net growth rate of sensitive cells under drug treatment. Similarly, the fraction of persister cells during the treatment is described by

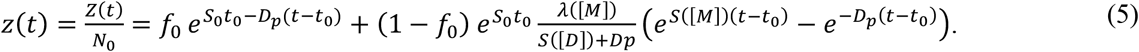

The quantity

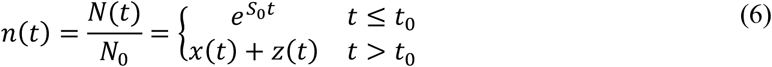

was used to “learn” the model and its parameters from the experimental data (see below).

#### Death rate of treated cells

The death rate of sensitive cells during drug treatment is assumed to increase from its basal level (*D*[0] ≡ *D*_0_) due to the effect of the drugs. In particular, based on previous evidence^32^, we model this dependence as

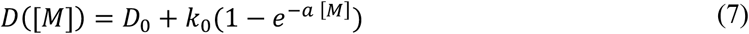

Where *k*_0_ is the maximum increase of death rate due to the drug effect. The parameter *a*^-1^ sets a characteristic drug concentration: for [*M*] << *a*^-1^ cells die with their unperturbed (basal) death rate *D*_0_, while for concentrations [*M*] >> *a*^-1^ the death rate reaches the maximal value *D*_0_ + *k*_0_. The empirical form Eq. (7) describes an exponential interpolation between these two extreme cases.

#### Transition rate to persistence

We considered four possible model variants for *λ*([*M*]): (i) a null model with no persistence state, corresponding to the case *λ* = 0 for any value of the drug concentration; (ii) a model with a drug independent transition rate *λ* = *λ*_0_; two models where the transition rate is linearly dependent on drug concentration, (iii) *λ*([*M*]) = *k*[*M*] and (iv) *λ*([*M*]) = *k*[*M*] + *λ*_0_. This latter functional dependence should be considered as the first term expansion of a nonlinear model with a rate saturating at high [*M*] values. The four model variants are associated to clearly distinguishable patterns as summarised in Extended Data Fig. 6.

#### Computer Simulations for the stochastic modeling

We simulated individual trajectories of the Markov process underlying the evolution of the MC-LD model. A well-known exact algorithm to simulate individual trajectories of a Markov process was provided by Gillespie^42^. However, this algorithm was too slow for the models and population sizes considered in this study. Therefore, we used a coarse-grained version of the algorithm, which groups together all stochastic events happening in discrete time intervals of fixed duration Δ*t*. This approximation is equivalent to assuming that events are independent within the time scale of Δ*t*. We adjusted the parameter Δ*t* so as to minimize the errors, while still keeping the simulations numerically feasible.

### Inference of parameters describing the cell population dynamics

#### Birth and death rates of sensitive cells

For the inference of the birth-death rates *b* and *d*, we used the data set described in the methods section *Growth rates of CRC clones before drug-treatment* (above), together with the model described in the methods section *Dynamics of sensitive (untreated) cells* (above). Our inference scheme is summarized in Extended Data Fig. 2a). The net growth rate *b-d* was estimated with an exponential fit against the observed growth curve of the two clones (WiDr and DiFi) in a time span of 4 days starting from 40×10^4^ cells/well in a 6-multiwell plate (Extended Data Fig. 2b). The value *d/b* was estimated from the observed asymptotic fraction of dead cells detected by flow cytometry analysis (Extended Data Fig. 2c). Finally, the values of the birth and death rate of sensitive cells were obtained by combining the two estimates (value of *b* - *d* and *d/b*). Estimated values are reported in Extended Data Table 1.

#### Calculation of growth curves from drug response assays

Growth curves of CRC clones under treatment, reported as fold-change of viable cells *vs* time of drug exposure, were calculated from the *doses-response assay* (Figs 1c and 2b) and from the *single-dose assay* (Figs 1d and 2c). We used the following strategy. Both data-sets consisted in measurements of cell viability (luminescence signal) with constant drug concentraction ([*M*]), evaluated at several time points (*t* = 1,2,.., *t_max_* days) and for a set of biological replicates (*i* = 0,1,.., *n*). We denote with 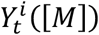 the value of cell viability of the *i^th^* biological replicate, of a drug-response growth assay performed with drug concentration [*M*], and evaluated at time *t*.

These ATP measurements present a day-to-day variability, which is reflected in a variability across replicates, but also present regularity in their behavior, which we aimed to extract. Specifically, the curves of different replicates look similar, but are affected by offsets that vary from day to day. Hence, we first normalized data corresponding to different biological replicates of the same clone in the following way. The values 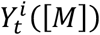 were multiplied by a normalization factor *c^i^*, the same for all the data collected in the same biological replicate, but different across replicates. These normalization factors were obtained by minimizing the total relative error across replicates 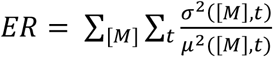, where 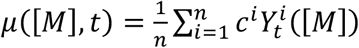 and 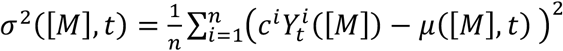 are the average and standard deviation of the viability normalized with weights *c^n^* across replicates, evaluated at time *t* and for drug concrentration [*M*].

Averages values and standard deviation computed with the optimal values of the normalization factors obtained by minimizing *ER* were then divided by the average viability measured at day 0 (*μ*([*M*], *t* = 0)) to obtain growth cuvers: y([*M*],*t*) =*μ*([*M*], *t*)/*μ*([*M*], *t* = 0) with associated standard deviation *σ_y_* ([*M*],*t*) = *σ*([*M*], *t*)/*μ*([*M*], *t* = 0).

#### TP model variants

Growth curves of WiDr and DiFi clones were used to infer model parameters of 8 (4 × 2) distinct model variants, i.e., configurations of the TP model with different assumptions of the two key model parameters: (i) the rate at which sensitive cells switch to persisters during the treament (*λ*) and (ii) the initial fraction of persister cells (*f*_0_). More in detail, we considered 4 model variants associated to the choice of the transition rate *λ*([*M*]) = {0,*λ*_0_, *k*[*M*],*k*[*M*] + *λ*_0_} which we combined with 2 variants using different ranges of the numerical values of the intial fraction of persisters (*f*_0_ = 0 and 0 < *f*_0_ < 1).

#### Inference of the TP model parameters

The parameters of the TP model (Eq. 6), with all the 4 model variants for the transition rate *λ*([*M*]) = {0, *λ*_0_, *k*[*M*], *k*[*M*] + *λ*_0_} and the two variants for the initial fraction of persisters (*f*_0_ = 0 and 0 < *f*_0_ < 1), have been inferred using a Bayesian framework. For the inference of TP model parameters in WiDr we used growth curves assessed from both the *doses-response assay* and the *single-dose assay*, while for DiFi we used only growth curves assessed from the *doses-response assay*. Posterior distributions of the model parameters were sampled using a Hamiltonian Monte Carlo (HMC) algorithm *(Python 3*, package *pymc3*, NUTS sampler)^43^. The likelihood function was set to the product of standard Gaussian likelihood functions over the observed data points, with parameters equal to the mean value and standard deviation of the data points. We assumed flat prior distributions of the model parameters; the corresponding supports, i.e. maximum and minimum values allowed, are reported in the Extended Data Table 2. For WiDr, model fit was performed including all the growth curves evaluated with dabrafenib concenteration [*M*] ≥ 0.041 *μM*, while for DiFi we included all growth curves evalutaed with cetuximab drug concenteration [*M*] ≥ 2.1 *nM*. Value of the model parameters describing the drug delay in the *single-dose assay* for DiFi (*t*_0_(*single*),*S*_0_(*single*)) were inferred with an indipendent model fit, while keepeing all the other parameters fixed (the remaining parameters were inferred from the growth curves derived from the *doses-response assay*). Values for these two parameters are reported in the Extended Data Table 2.

#### Comparison between TP model variants

The logic flow of the comparison between model variants within our inference scheme is summarized in Extended Data Fig. 6. The eight TP model variants (2 choices of *f*_0_ for each of the 4 *λ* models) were compared by means of the standard Bayesian Information Criterion (BIC) and the Akaike Information Criterion (AIC) (Extended Data Table 3). These quantities measure model performance keeping into account the number of parameters used (penalizing model variants with more parameters). We found that the best *λ* model the for WiDr is the one where the transition rate to persistence is linearly proportional to the drug concentration *λ*([*M*]) = *k*[*M*], while for the DiFi the data is best described by a TP model variant with a constant transition rate *λ*([*M*]) = *λ*_0_ (Extended Data Table 3 and Extended Data Fig. 6). Additionally, we found that both the AIC and BIC indicate that the drug-induced scenario (*f*_0_ = 0) is the preferred variant for all the cell lines (Extended Data Table 3 and Supporting Fig. 6).

#### Choice of the TP model variant for the inference of the mutation rate

The parameters used to calculate the mutation rates were inferred using the following TP model variants: (i) transition rate to persistence linearly proportional to the drug concentration *λ*([*M*]) = *k*[*M*] and 0 < *f*_0_ < 1 for WiDr cells and (ii) constant transition rate *λ*([*M*]) = *λ*_0_ and 0 < *f*_0_ < 1 for DiFi cells. For both clones, the choice of the best *λ* model was informed by the BIC and AIC indices (Extended Data Table 3), while in both cases we made use of the variant with 0 < *f*_0_ < 1 even though *f*_0_ = 0 was preferred by the BIC and AIC indices (Extended Data Table 3). This was done in order to have a realistic estimate of the maximum value of *f*_0_ which is still compatible with the experimental growth curves data. In this way we were able to infer the value of the mutation rate taking into account the effect of a fraction of persisters that could have been present in the population before drug administration. The fit of these models to the experimental data is shown in Fig. 2 and Extended Data Fig. 4, while the statistics of the corresponding model parameters are reported in Extended Data Table 4. Corresponding posterior distributions are shown in Extended Data Fig. 5. Plots of the posterior distributions were obtained with the software GetDist^44^. Note that the approach used in Figs 2d and 4d and discussed in the main text does not focus only on the optimal model variant (i.e., *f*_0_ = 0), but explores all the possible values of *f*_0_.

#### Distribution of persister cell abundance

The TP model presented in the previous section was extended to include stochastic effects to investigate the distribution across wells of the number of persister cells at a given time point (well-to-well number variability). To this aim, we made use of computer simulations (see methods section *Computer Simulations for the stochastic modeling).* We found that in the drug-induced scenario, i.e., in the absence of persister cells before drug treatment (*f*_0_ = 0), and for time of drug treatment » 1 *day*, the number of cells is distributed according to a Poisson distribution (see Extended Data Fig. 7). Extended Data Fig. 7b exploits this relation to verify that the drug-induced scenario is compatible with the observed dynamics of both CRC cell clones, using the experimentally measured well-to-well distribution of cell numbers measured after 3 weeks of treatment. The cumulative distribution of the cell viability measurements across well of drug-tolerant persisters cells is indeed compatible with a Poisson cumulative distribution (Extended Data Fig. 7b). Since the ATP assay measures the number of cells through an unknown constant of proportionality between the ATP content ad the actual number of viable cells, in order to fit the observed distribution of ATP assay readouts to a Poisson distribution we proceeded as follows. We computed the value 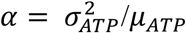, i.e., the ratio of the variance 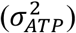 over the mean (*μ_ATP_*) of the ATP measurements across the wells. The best fit to the distribution of the ATP measurements was the function 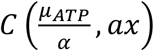, where *C*(*μ,x*) is the cumulative distribution function of a variable (x) distributed according to a Poisson distribution with mean *μ*.

### Inference of the mutation rate from a two step fluctuation assay

#### Probability for the emergence of at least one mutant

This section derives from the MC-LD model an approximate expression for the probability of the emergence of one mutant in an expanding population of cells in a given time interval [0,*T*]. In the MC-LD model, we denote with N(*t*) the number of viable cells present in the population at time *t*, and with *μ* the effective rate at which one individual becomes a mutant. We assumed that mutant individuals in the population have the same dynamical rates as untreated cells, i.e., they divide with rate *b* and die with rate *d.* Because of reproductive fluctuations (genetic drift), cells carrying drug-resistance mutations can still go extinct, and only a fraction of the mutants will “establish” in the population, and survive. We refer to these cells in the following as “established mutants”, using the standard terminology of population genetics. The probability of surviving stochastic drift in a time interval Δ*t* is a well-known result of the birthdeath process^37,38^, which reads

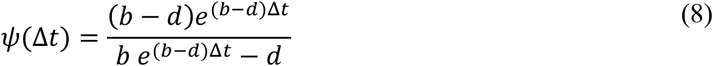

With these assumptions and up to the first order in *μ* (i.e., assuming a sufficiently small mutation rate) the expected number of emerging mutants establishing in [0,*T*] is given by

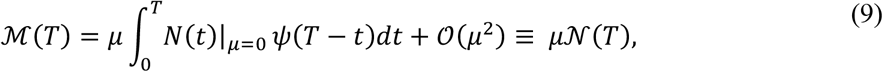

where we have defined 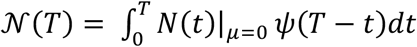. The number of established mutant cells is Poisson distributed with expected value 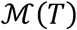 and consequently the probability of having at least one mutant is given by

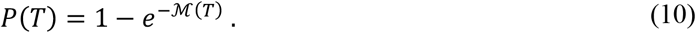

In the context of the fluctuation test, this probability is estimated by 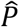, i.e., the fraction of wells that have developed resistant mutants by the time *T* after treatment. The generalized estimator of the mutation rate then takes the form

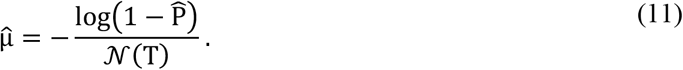

This general form of the estimator is valid for both pre-existing and persisters-derived resistant cells, and has been tested with synthetic data (see Fig. 4c). More specific expressions that can be used with data are found below.

#### Estimator of mutation rate for sensitive cells

As in a standard fluctuation test, to estimate the spontaneous mutation rate of sensitive (untreated) cells *μ_s_*, we measure the fraction 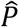 of wells that developed resistance before treatment initiation (i.e., during the expansion phase in absence of drug). Since the latters grow unperturbed when treatment is applied, the early-emerging resistant clones, arising within the first 3-4 weeks of drug exposure in our experimental setting represent the cells that developed resistance before treatment (see Extended Data Fig. 9) The first part of the MC-LD model uses an estimator for the mutation rate of sensitive (untreated) cells from the number of pre-existing mutants that were generated before the exposure to the drug. Following the approximations found in Eq (11), for this case we have

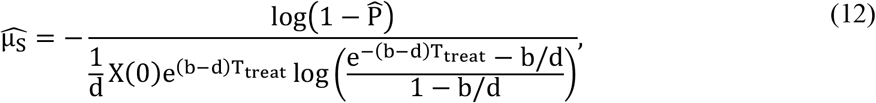

where *X*(0) is the number of sensitive cells present in the population at the beginning of the fluctuation test. As described in the main text, during our fluctuation test sensitive cells were allowed to expand for a time *T_treat_* before starting the treatment. The standard deviation of the estimated value of 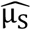 was obtained by error propagation of the standard deviation of the fraction of observed wells harboring resistant clones 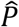, and is defined by the following expression^45^

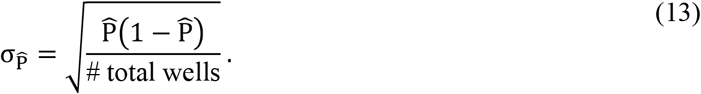

The mutation rate Eq. (12) is a *chronological* rate, i.e. it quantifies the number of resistant cells emerging in the population per unit of time and per individual. As in this case new sensitive cells are generated by cell division, this rate can be converted into units of generation 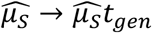, where *t_gen_* = 1/*b* is the duration of one generation. The values of the observed number of wells with preexisting colonies of resistant cells are show in Fig. 3e, while the values of the inferred mutation rate are reported in Extended Data Table 5.

#### Estimator of mutation rate for persister cells

To estimate the mutation rate of persister cells *μ_p_* in the second part of the MC-LD assay, we extended the experimental procedure described for sensitive cells, prolonging the treatment of persisters-containing wells and therefore performing a fluctuation assay which evaluates the fraction of the remaining wells developing resistant clones. As described above (Eq. 10), the expected fraction of resistant cells derived from persisters cells and evaluation at time *T* is given by

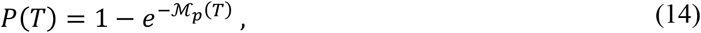

where

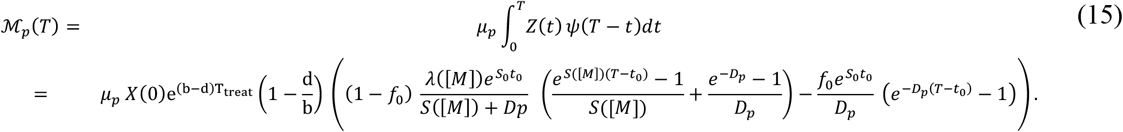

where 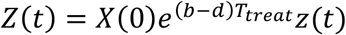, is the number of persister cells expected at time *t* (product of the expected fraction of persister cells *z*(*t*) from Eq.(5) and the total number of cells present in the wells at the beginning of treatment *X*(0)*e^(b-d)T_treat_^*), and we have used the approximation 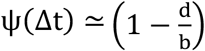. The estimator for the mutation rate of persister cells can be obtained by matching the expected probability Eq (14) to the observed fraction of wells with growing colonies observed 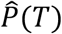:

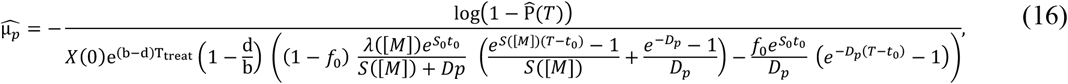

Our analitcal derivation (Eq.s 14, 15 and 16) was validated with simulations of the MC-LD model, by testing the estimator (Eq. 16) with synthetic data (see Fig. 4c).

#### Bayesian inference of the mutation rate of persister cells

The mutation rate of persister cells was inferred with a Bayesian framework, in order to account for the uncertainty of the value of the initial fraction of persister cells, *f*_0_. The bayesian inference of *μ_p_* was obtained by fitting the MC-LD model expectation (Eq.s 14 and 15) to the observed fraction of wells with late-emerging resistant clones (evaluated at T=11 weeks for WiDr and at T=15 weeks for DiFi). The likelihood function was set to a standard Gaussian likelihood with mean equal to 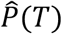 and standard devation 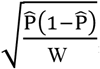, where *W* is the number of wells that did not harbour early emerging resistant clones. The model expectation was computed using the model parameters inferred for the TP model (Extended Data Table 4). For the inference of the mutation rate of DiFi B6, we used a value *t*_0_ = 0, consistently with the data-set of Extended Data Fig. 4. We inferred two parameters: (i) the initial fraction of persister cells *f*_0_ and (ii) the fold increase of the mutation rate of peristers cells vs sensitive cells 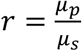, i.e., the mutation rate was inferred a the product of this parameter and the inferred value of the mutation rate of sensitive cells, *μ_s_* (see above). For the first parameter, *f*_0_, we used an exponential prior distribution with the same mean as the one reported in Extended Data Table 4, i.e., we imposed a prior on this parameter which is equal to the posterior distribution obtained when fitting the TP model to the growth curve data (Fig. 2d). For the second parameter, 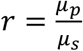 we used a flat distribution with minimum value 1 and maximum value 500. The posterior distributions for *f*_0_ and for the fold increase of the mutation rate (shown in Fig. 3d) were sampled using a Hamiltonian Monte Carlo (HMC) algorithm *(Python 3*, package *pymc3*, NUTS sampler)^43^. Values of the estimated mutation rate and its standard deviation, reported in Extended Data Table 5, were computed as *μ_p_* = < *r* > *μ_s_*, and *σ_μ_*, = *σ_r_ μ_s_*, where < *r* > and *σ_r_* are the mean and the standard deviation of the posterior distribution of the parameter *r*.

## Materials availability

The CRC cell clones generated in this study are available through Alberto Bardelli (Department of Oncology, University of Torino) under a Material Transfer Agreement.

## Data and code availability

Data and codes used for the analysis will be available as a repository on Mendeley Data (doi:10.17632/mvfm7hs9kw.1).

## Acknowledgments

We are grateful to A. Ciliberto, M. Osella, L. Trusolino and A. Amir for useful discussions and critical reading of the manuscript. This work was supported by FONDAZIONE AIRC under 5 per Mille 2018 - ID. 21091 program - P.I. Bardelli Alberto; Group leaders Di Nicolantonio Federica and Bertotti Andrea; AIRC, Associazione Italiana per la Ricerca sul Cancro AIRC-IG (REF: 23258) - P.I. Marco Cosentino Lagomarsino; AIRC IG 2017 N° 20697 - P.I. Andrea Bertotti; European Research Council Consolidator Grant 724748, BEAT - PI Andrea Bertotti; H2020 grant agreement no. 635342-2 MoTriColor to A.Bardelli; AIRC under IG 2018 - ID. 21923 project - P.I. Bardelli Alberto; AIRC-CRUK-FC AECC Accelerator Award contract 22795 to A.Bardelli; Ministero Salute, RC 2020 to A.Bardelli; Fondazione Piemontese per la Ricerca sul Cancro-ONLUS 5 per mille 2015 Ministero della Salute to A.Bardelli and F.D.N. Simone Pompei was supported by Fondazione Umberto Veronesi.

## Author contributions

M.R., A.Bardelli and M.C.L. conceived the study and contributed with key ideas at different stages. M.R. and A.S. performed the biological experiments. S.P., M.C., M.G. and M.C.L. conceived the modeling framework. S.P., M.C. and M.G. performed data analysis, model simulations, and analytical calculations. G.C., A.Bertotti and F.D.N. contributed to data discussion. M.R., S.P., A.Bardelli and M.C.L. wrote the paper. All authors read and approved the final version of the paper. M.R., S.P., A.S. and M.C. equally contributed to the study. A.Bardelli and M.C.L. jointly supervised the study.

## Competing interests

A.Bardelli reports receiving commercial research grants from Neophore; is an advisory board member/unpaid consultant for Roche, Illumina, Guardant, and Third Rock; holds ownership interest (including patents) in Neophore and Phoremost; and is an advisory board member/unpaid consultant for Horizon Discovery, Biocartis, and Neophore. All the other authors declare no competing interests.

## Extended Data Figures

**Extended Data Fig. 1.**
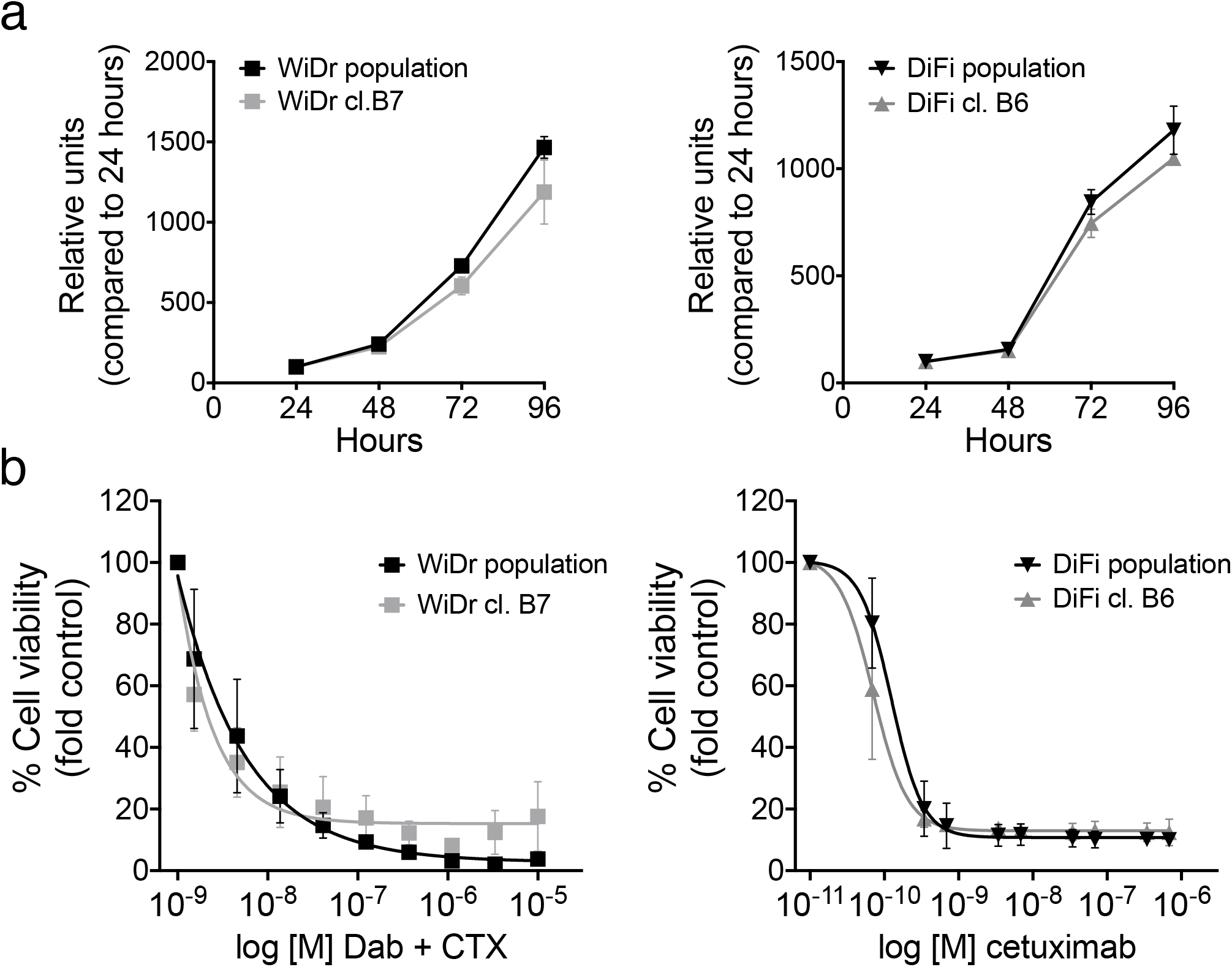
Growth kinetics of CRC cell populations and individual clones. **a**, The growth kinetics of WiDr and DiFi clones were compared with that of parental population at the indicated timepoints. Cell viability was assayed by the ATP assay. **b**, The indicated cells were treated with increasing concentrations of dabrafenib (Dab) + 50μg/mL cetuximab (CTX) (WiDr) and increasing concentrations of cetuximab (DiFi). Cell viability was measured with the ATP assay after 5 (WiDr) or 6 days (DiFi). Results represent the average ± SD (n=3).

**Extended Data Fig. 2.**
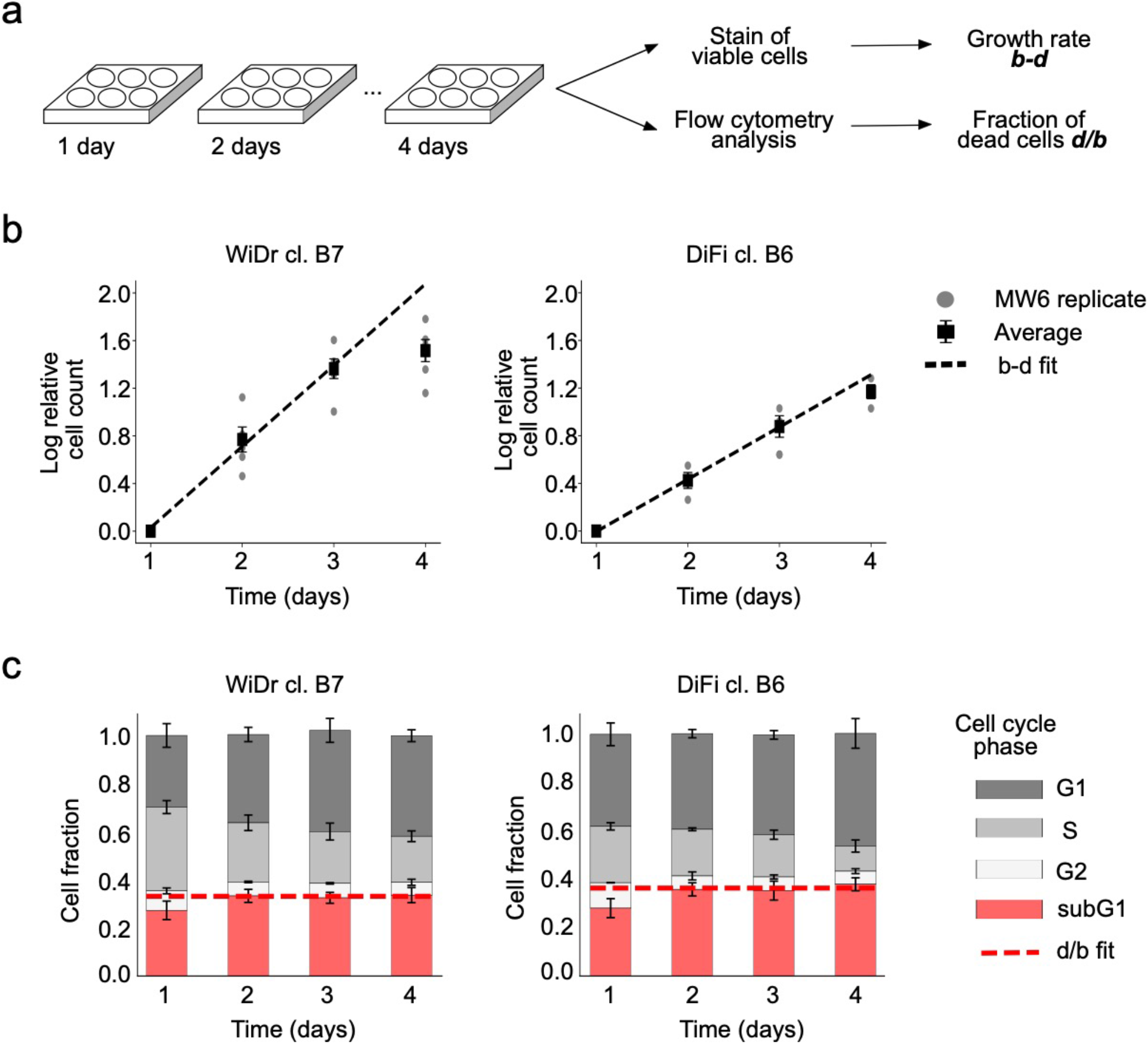
Birth and death rates of CRC cell clones. **a**, Schematic representation of the experimental setting used to evaluate birth and death rates of CRC cells. **b**, To establish the growth rate *(b-d)*, the indicated cell models were seeded in multiple 6-multiwell plates (MW6) at ~ 4×10^5^ cells/well, and the number of viable cells was measured by manual count using trypan blue staining at the indicated time points. Grey dots represent individual biological replicates, each reported as mean of two technical replicates. Black squares represent average of biological replicates reported as mean ± SD (n=5 for WIDr, n=3 for DiFi). The black dashed line shows the best exponential fit (here represented as a linear fit on the log number of relative cell count). **c**, Cell cycle distribution of CRC clones measured by propidium iodide staining and flow cytometry analysis at the indicated time points. The fraction of cells in sub-G1 phase was used to estimate the death rate (*d/b*). Bars represent mean ± SD (n=3). Red dashed line shows the best fit for the value *d/b*, which is the expected asymptotic value of the fraction of dead cells δ.

**Extended Data Fig. 3.**
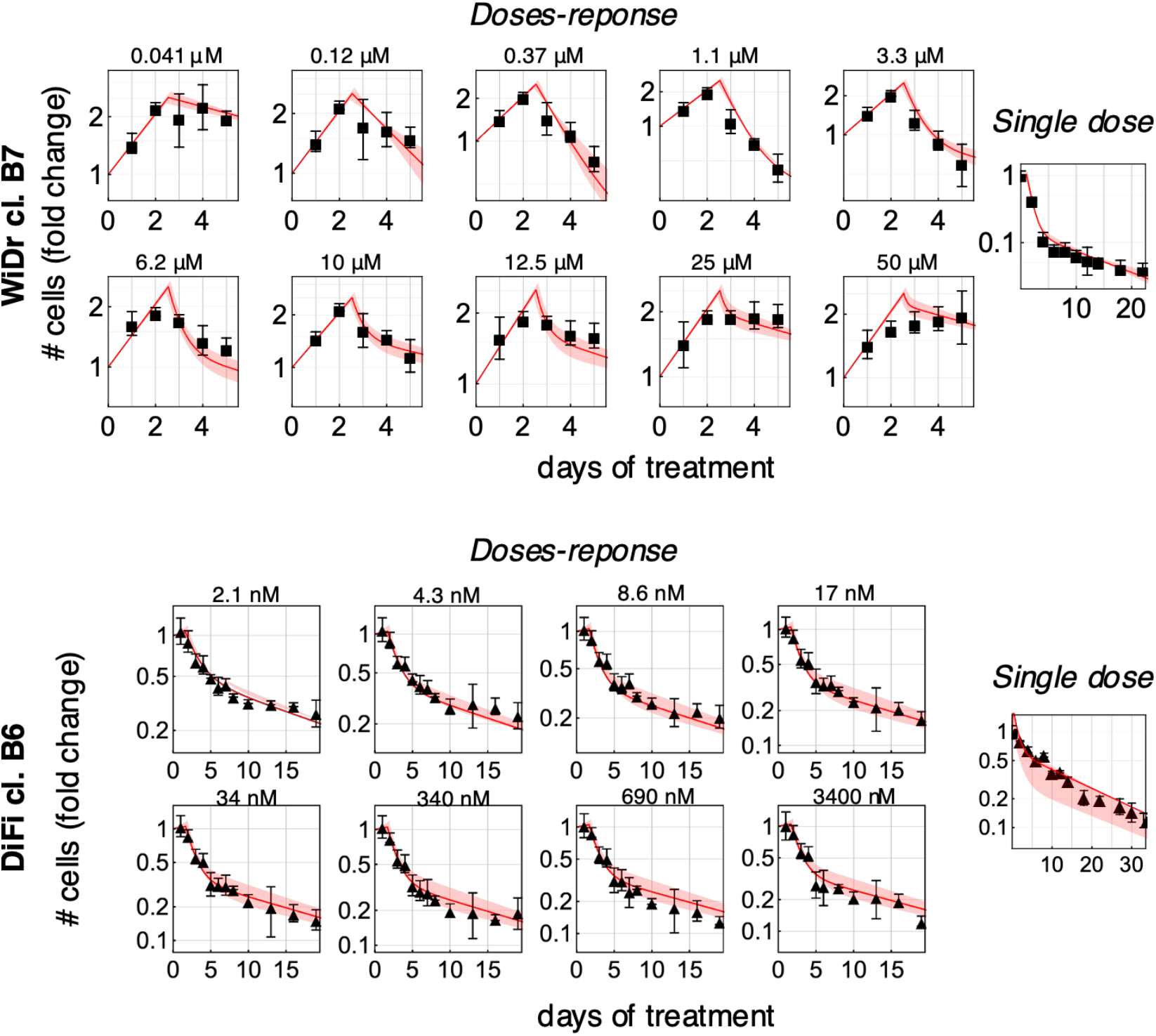
Transition to persisters (TP) model of CRC cells. Fitted experimental data of the *doses-response* (n=3 for WiDr, n=5 for DiFi, left side) and *single-dose* (n=2, right side) datasets are indicated with black dots and error bars; red lines and shadowed areas represents the model fit (Credible Interval [2.5, 97.5]%). The drug concentration is specified above each plot.

**Extended Data Fig. 4.**
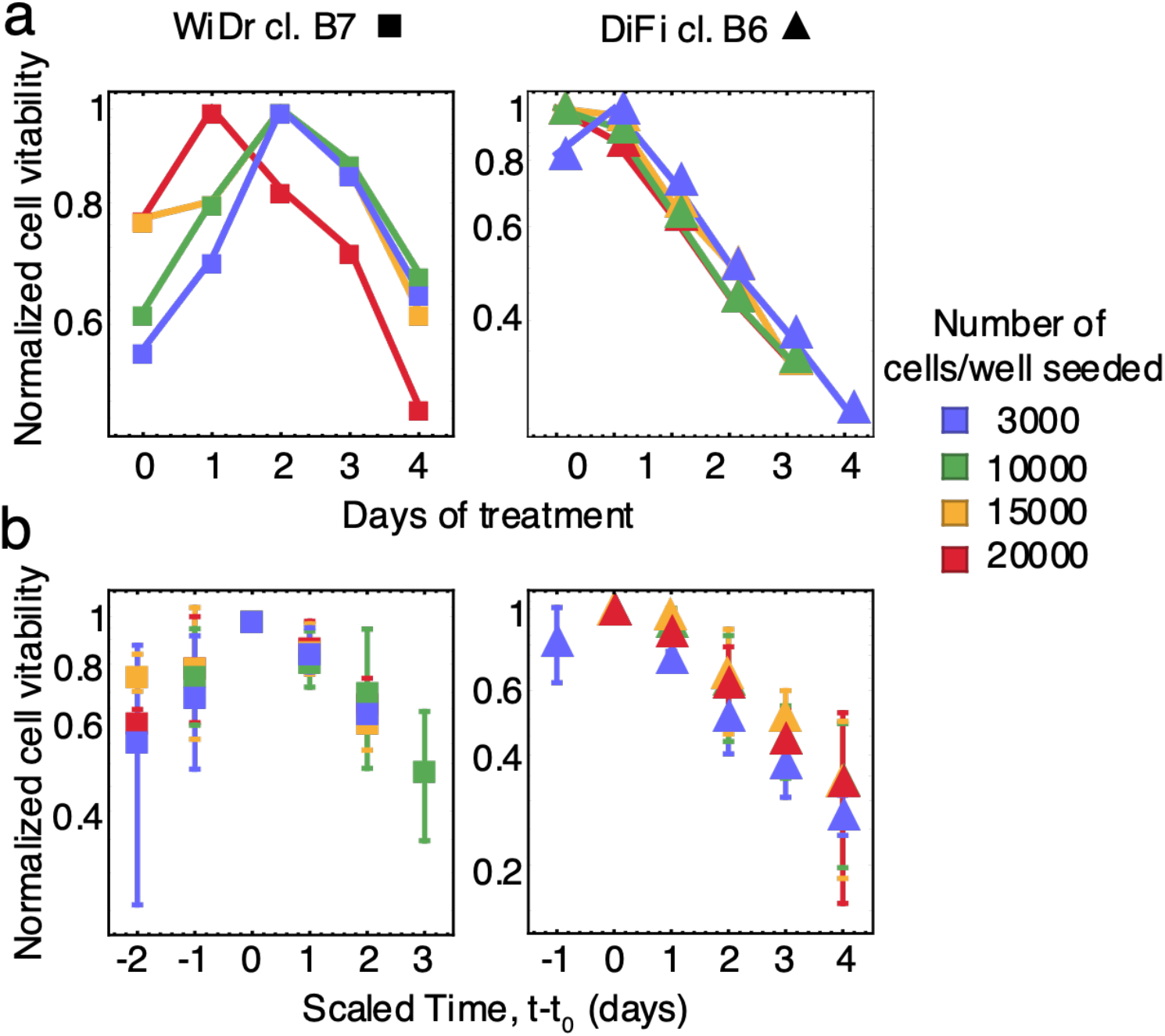
Impact of seeding density on cell growth dynamics. **a**, Cells seeded at indicated densities, were treated with 1μM dabrafenib + 50μg/ml cetuximab (WiDr) or 100 μg/ml cetuximab (DiFi). Cell viability was measured with the ATP assay at indicated time points. **b**, Cell growth assays performed with different initial number of cells display the same dynamics after an initial delayed effect of the treatment. All the values were normalized to the maximum value of cell viability measured. Time values from (**a**) were scaled to *t*_0_ (time of delayed drug effect) indicating the time when the maximum cell viability was reached.

**Extended Data Fig. 5.**
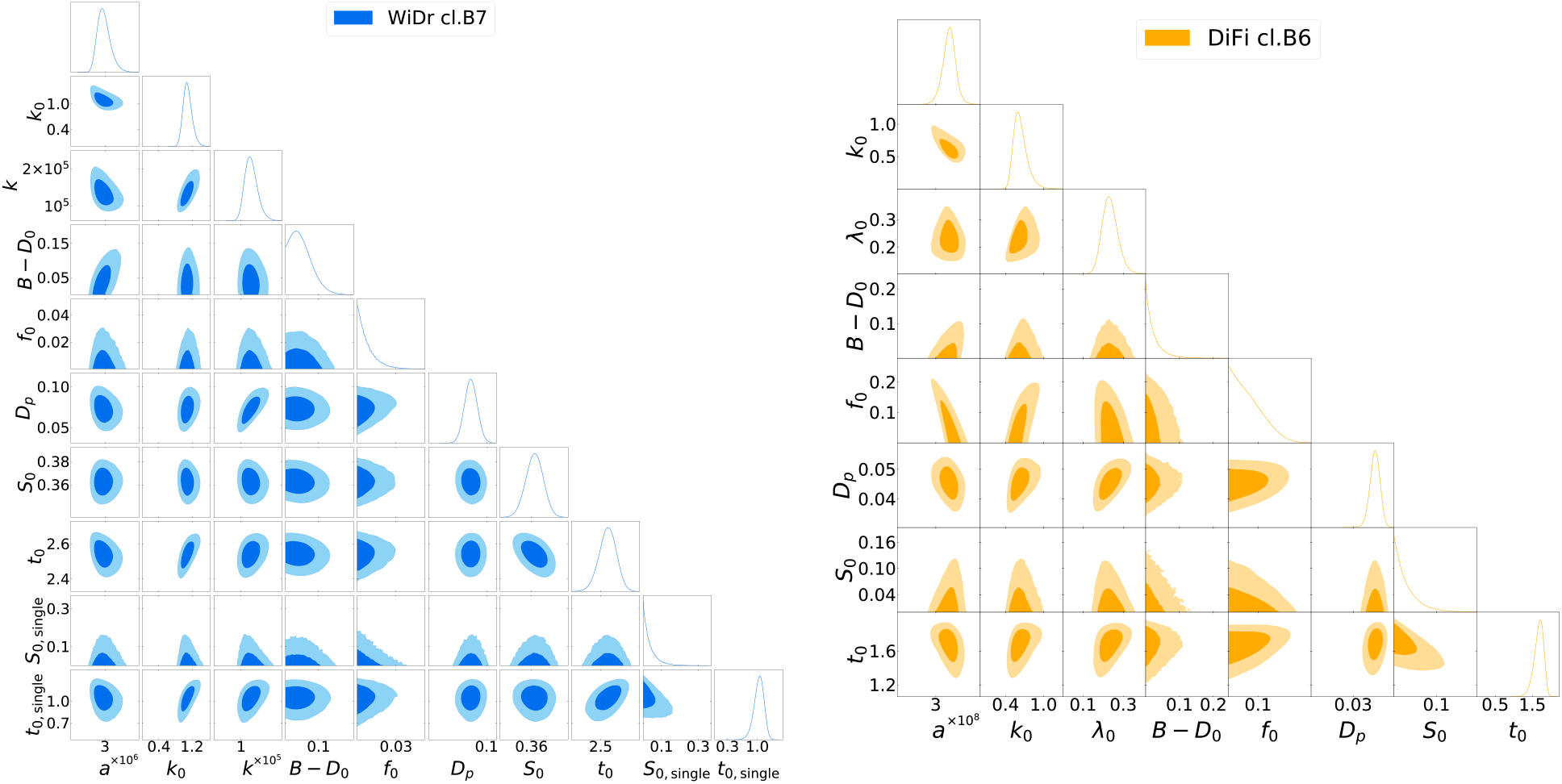
Posterior distributions of the inferred parameters for the transition to persister (TP) model. For each model we show posterior distributions of the inferred parameters. In the diagonal we show distributions marginalized to one parameter, off-diagonal plots show contour plots at one and two sigmas (dark and light orange, respectively) of distributions marginalized to two parameters. See Methods for description of Bayesian inference. The model parameters are the following: ***a***: inverse of the characteristic drug concentration. For a drug concentration [*M*]<<*a*^-1^ cells die with their unperturbed death rate *D*_0_, while for concentrations [*M*]>>*a*^-1^ the death rate *κ* reaches the maximal value (*D*_0_ + *k*_0_). ***k**_0_*: maximum death rate due to the drug. ***k*** coefficient of proportionality between drug concentration and rate of transition of sensitive cells to persister cells (*λ*=*k*_0_[*M*]). ***t_0_:*** time of the delay of the drug effect after administration. ***B-D_0_:*** growth rate in absence of drug (birthrate (*B*) minus death rate *D_0_*). ***S_0_:*** growth rate observed in the temporal window between drug administration and drug effect. ***λ_0:_*** rate of transition of sensitive cells to persister state. ***D_p_:*** death rate of persister cells. ***t_0 (single):_*** delay in time of drug effect after treatment administration in the *single-dose assay. **S_0(single):_*** observed growth rate in the time window between treatment administration and drug effect (t < ***t_0(single)_***) in the *single-dose assay*.

**Extended Data Fig. 6.**
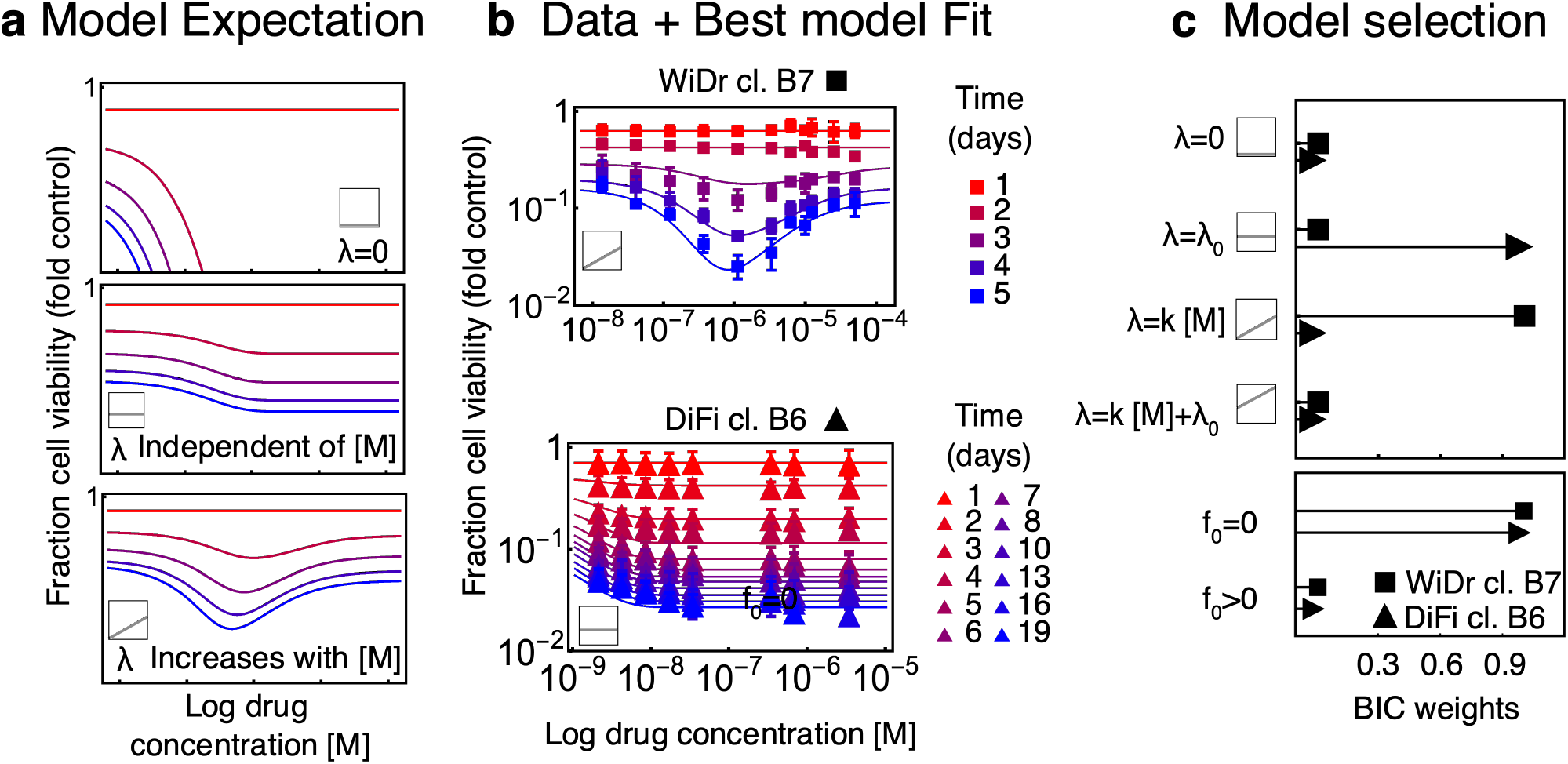
Bayesian inference of experimental data defines a model for the transition to persistence. **a**, Schematic representation of the expected pattern of the *doses-response* curves for the three different model configurations: (i) null model with no transition to persistence (λ = 0, top), (ii) model with constant transition rate (mid), (iii) model with transition rate proportional to the drug concentration (bottom). Square plots indicate the dependence of λ vs [M]. **b**, Experimentally measured dose-responses growth curves and best model fit. The *doses-response* datasets were normalized to the growth of the untreated cells using the best model parameters (Extended Data Table 4). **c**, In the top panel, we show the Bayesian weights to compare the 4 λ models. Bayes weights are defined as 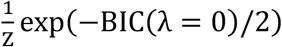 for the model as 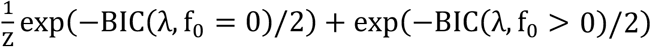 for the other models. The partition function Z ensures the global normalization Z = ∑_λ,f_0__ exp(-BIC(λ, f_0_)/2). Similarly, in the bottom panel we show the two Bayes factors to compare model configuration with 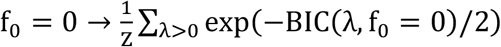 (drug induced scenario) and for 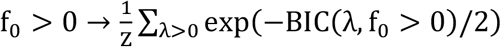. Values of the BIC values are reported in Extended Data Table 3.

**Extended Data Fig. 7.**
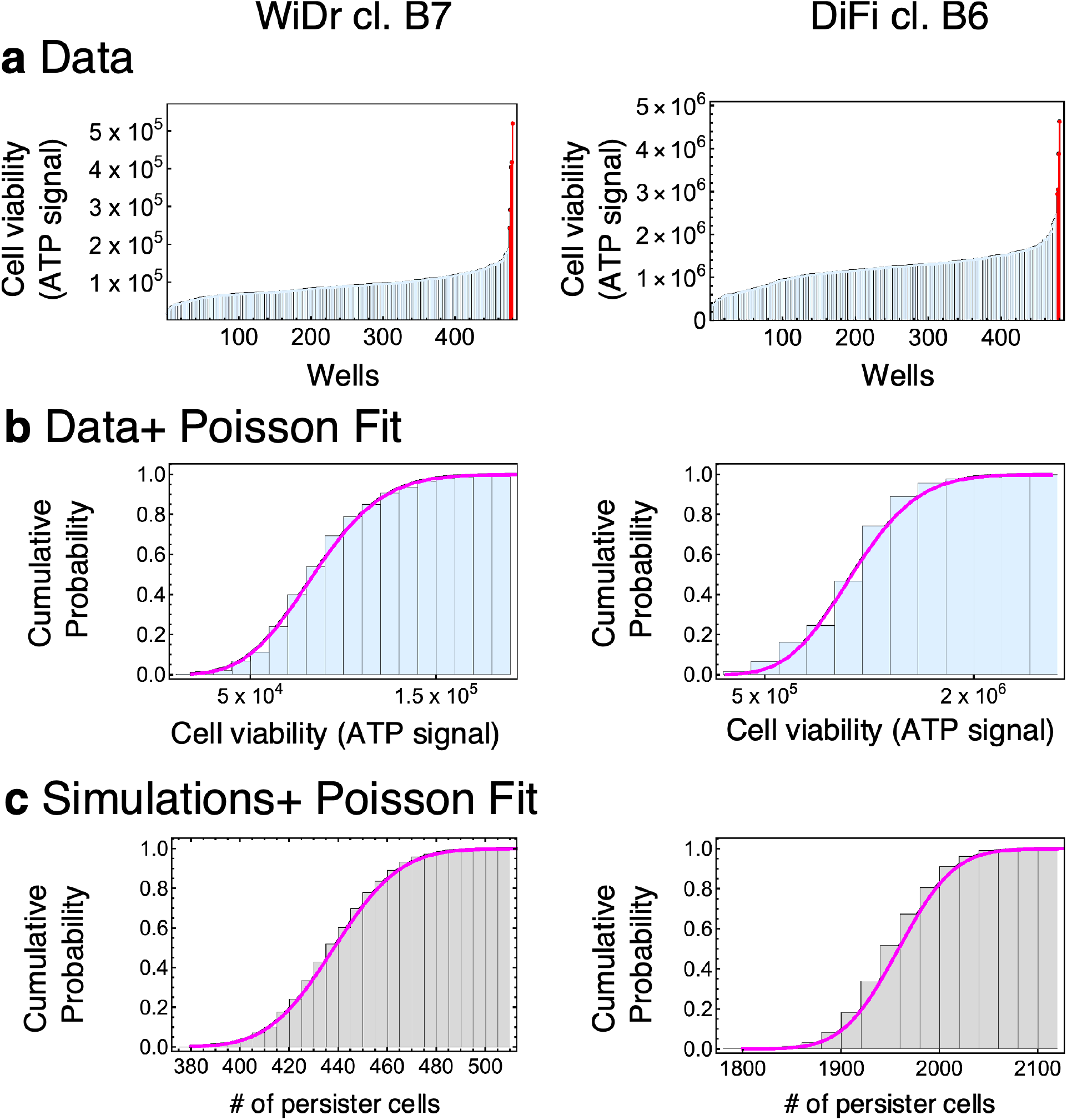
Distribution of persisters abundance across wells is compatible with the TP model expectation of *f*_0_=0. **a**, CRC clones were seeded in multiple 96-multiwell plates and allowed to expand for about 8 cell divisions in the absence of drug, and then treated with 1μM dabrafenib + 50μg/ml cetuximab (WiDr) or 100 μg/ml cetuximab (DiFi) for 3 weeks. Cell viability was determined by ATP assay. Wells containing rapidly proliferating pre-existing resistant colonies are marked in red, while the remaining wells contained a small number of viable cells, which we identified as drug-tolerant persisters (indicated in light-blue). Each bar represents one well. One representative experiment of two independent replicates is reported. **b**, Cumulative distribution of persisters cell viability across wells (light-blue bars) is compatible with a Poisson cumulative distribution (magenta solid line, see methods for details on the fitting procedure). One representative experiment of two independent replicates is reported. **c**, The simulation of a stochastic model for the transition to persistence shows that in the case of *f*_0_ = 0 (i.e., in the drug-induced scenario) the number of persister cells per well is expected to have a Poisson distribution. Simulations were performed using the TP model fit values as input parameters, and setting the initial population size to 15000 cells/well. Simulations were stopped at 21days; the distribution was evaluated using 1000 independent simulations.

**Extended Data Fig. 8.**
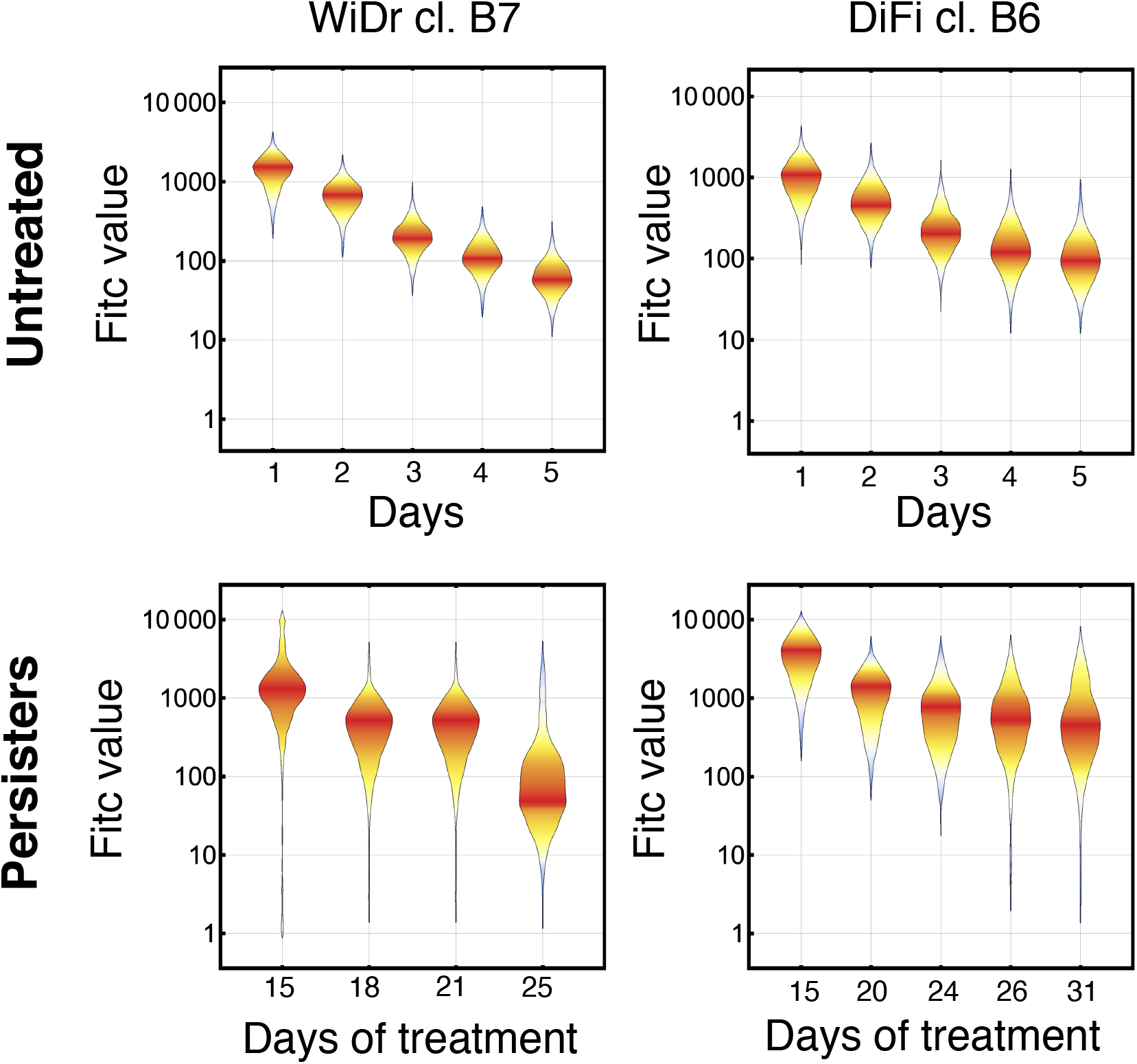
A fraction of persisters cells slowly replicate during drug treatment. Distribution of the CFSE signal (Fitc) measured by flow cytometry is reported for the indicated timepoints. WiDr and DiFi cells were grown in standard conditions (untreated) or treated with 1μM dabrafenib + 50μg/ml cetuximab or 100μg/ml cetuximab, respectively, for 2 weeks until the emergence of surviving persister cells. Both untreated and persister cells were stained with CFSE to quantify cell division and fluorescent signal (Fitc) was analyzed by flow cytometry at indicated time points. One representative experiment (n=2 untreated cells; n=3 persisters) is reported.

**Extended Data Fig. 9.**
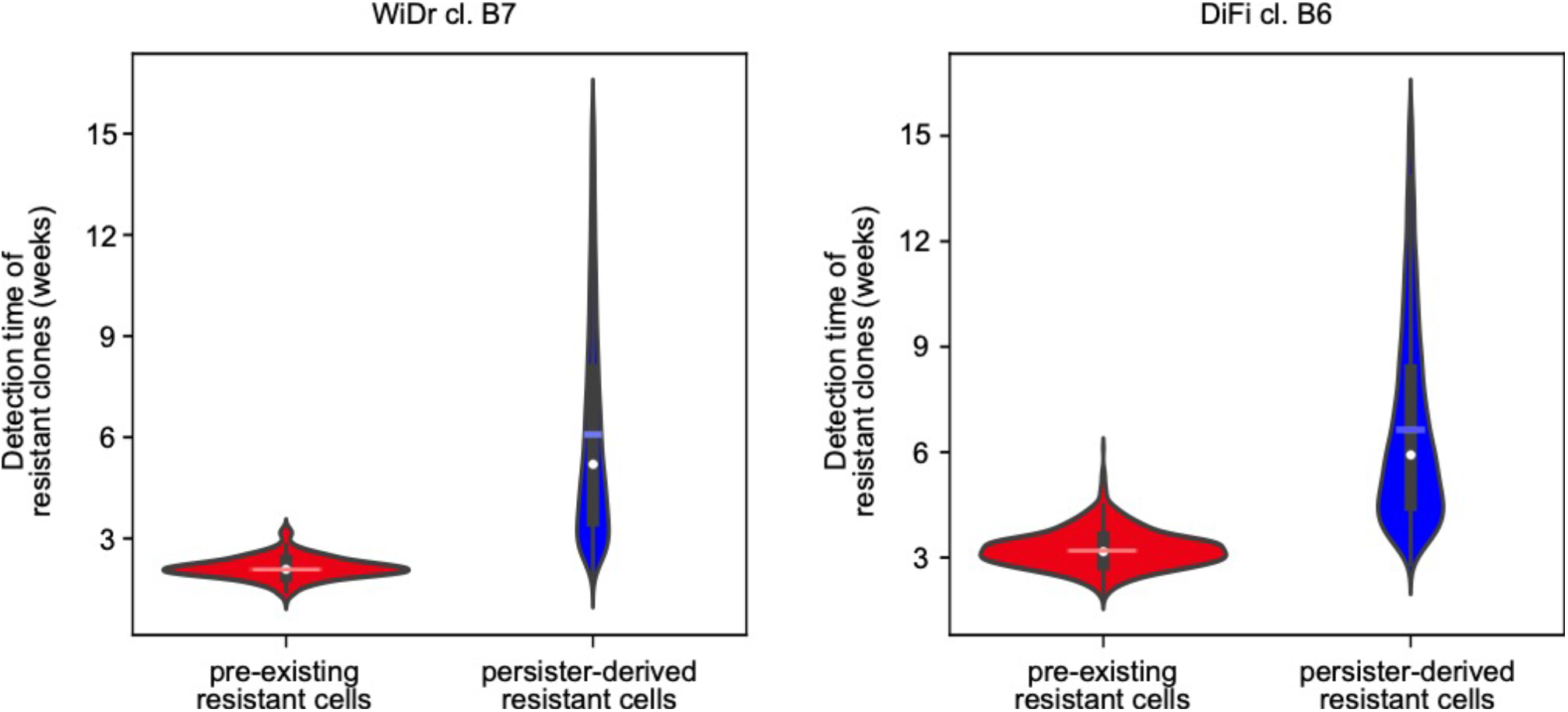
Pre-existing and persister-derived resistant cells emerge at different time points according to the MC-LD model. For each clone we simulated the MC-LD fluctuation test using experimentally determined cellular parameters. The violin plot showing the time of emergence distribution is reported for pre-existing resistant cells (indicated in red) and persisters-derived resistant cells (depicted in blue). Median and mean of the distribution are represented as a white dot and a nuanced horizontal line, respectively. The black vertical line indicates the interval corresponding to the first and third quartile of the distribution. As previously reported, after about 3-4 weeks the vast majority of early-emerging resistant clones originate from pre-existing resistant cells, while persisters-derived resistant cells slowly accumulate subsequently over time. We run 50 simulations using the best model fit as input parameters. In each simulation, we simulated 1920 in silico independent wells. A well was considered to harbor a resistant clone when the number of resistant cells was above 20,000 units.

## Extended Data Tables

**Extended Data Table 1.** Growth parameters of CRC cells in the absence of drug treatment. The table lists experimentally assessed birth and death rates for the indicated CRC cell clones. Mean values and standard errors are reported.

|  | WiDr cl. B7 |  | DiFi cl. B6 |  |
| --- | --- | --- | --- | --- |
|  | Value | S.E. | Value | S.E. |
| Birth rate (b) ($days^{-1}$) | 1.02 | 0.08 | 0.7 | 0.02 |
| Death rate (d) ($days^{-1}$) | 0.34 | 0.09 | 0.26 | 0.02 |

**Extended Data Table 2.** Prior distributions for persister-transition model fit parameters. We assumed flat prior distributions for each of the TP-model parameters. The tables list the min and max allowed values as support of the indicated distributions.

| WiDr cl. B7. |  |  |
| --- | --- | --- |
| Parameter (units) | Min value | Max value |
| $a$ ($M^{-1}$) | $10^5$ | $10^9$ |
| $k_0$ ($days^{-1}$) | 0 | 6 |
| $k$ ($M^{-1}days^{-1}$) | 1 | $10^6$ |
| $B - D_0$ ($days^{-1}$) | 0 | 1 |
| $f_0$ | 0 | 1 |
| $D_p$ ($days^{-1}$) | 0 | 0.2 |
| $S_0$ ($days^{-1}$) | 0 | 1 |
| $t_0$ ($days^{-1}$) | 2 | 3 |
| $t_0(Single)$ ($days^{-1}$) | 0 | 3 |
| $S_0(Single)$ ($days^{-1}$) | 0 | 1 |

| DiFi cl. B6. |  |  |
| --- | --- | --- |
| Parameter (units) | Min value | Max value |
| $a$ ($M^{-1}$) | $10^7$ | $10^9$ |
| $k_0$ ($days^{-1}$) | 0 | 3 |
| $\lambda_0$ ($days^{-1}$) | 0 | 1 |
| $B - D_0$ ($days^{-1}$) | 0 | 1 |
| $f_0$ | 0 | 1 |
| $D_p$ ($days^{-1}$) | 0 | 0.2 |
| $S_0$ ($days^{-1}$) | 0 | 1 |
| $t_0$ ($days^{-1}$) | 1 | 2 |
| $t_0(Single)$ ($days^{-1}$) | 0 | 2 |
| $S_0(Single)$ ($days^{-1}$) | 0 | 5 |

**Extended Data Table 3.**
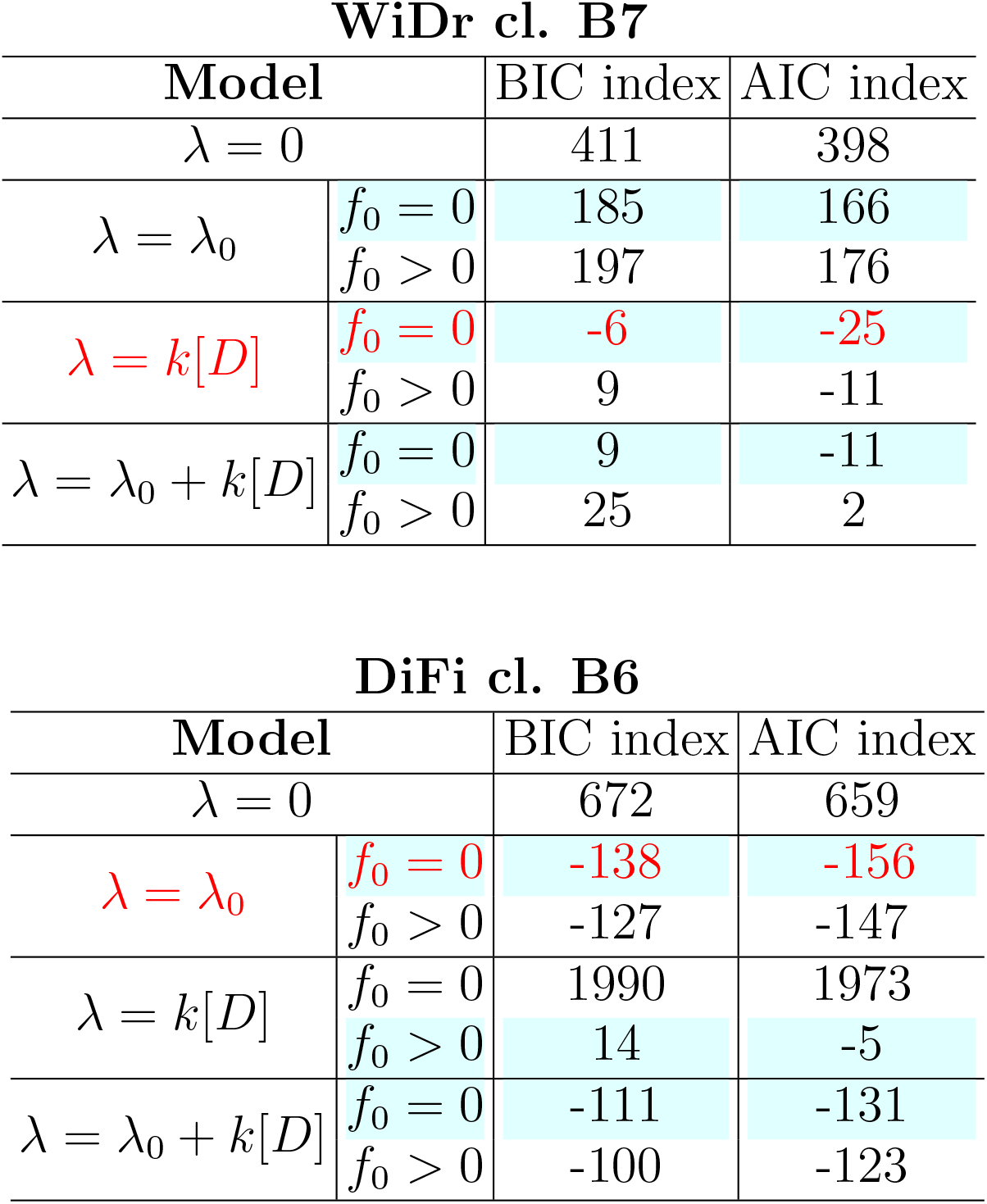
Model selection for the transition to persister (TP) model variants. The table lists the Bayesian Information Criterion (BIC) and Akaike Information Criterion (AIC) of all the transition to perister (TP) model variants. Lower AIC and BIC values correspond to better model performance, accounting for the different number of parameters. 4×2 models were inferred (2 choices of *f*_0_ for each of the 4 λ models) for the indicated clones. For each transition rate (λ), the model *f*_0_ (indicating the fraction of persister cells before treatment initiation) with the relative minimum value of both AIC and BIC index is highlighted in cyan; while the model with the global minimal value of the indices, i.e., the best TP model fit, is indicated in red. The λ models are recapitulated in Extended Data Fig. 6.

**WiDr cl. B7**
| Model |  | BIC index | AIC index |
| --- | --- | --- | --- |
| $\lambda = 0$ | | 411 | 398 |
| $\lambda = \lambda_0$ | $f_0 = 0$ | 185 | 166 |
| | $f_0 > 0$ | 197 | 176 |
| $\lambda = k[D]$ | $f_0 = 0$ | -6 | -25 |
| | $f_0 > 0$ | 9 | -11 |
| $\lambda = \lambda_0 + k[D]$ | $f_0 = 0$ | 9 | -11 |
| | $f_0 > 0$ | 25 | 2 |

**Extended Data Table 3. Model selection for the transition to persister (TP) model variants.**
| Model |  | BIC index | AIC index |
| --- | --- | --- | --- |
| $\lambda = 0$ | | 672 | 659 |
| $\lambda = \lambda_0$ | $f_0 = 0$ | -138 | -156 |
| | $f_0 > 0$ | -127 | -147 |
| $\lambda = k[D]$ | $f_0 = 0$ | 1990 | 1973 |
| | $f_0 > 0$ | 14 | -5 |
| $\lambda = \lambda_0 + k[D]$ | $f_0 = 0$ | -111 | -131 |
| | $f_0 > 0$ | -100 | -123 |

WiDr cl. B7

**Extended Data Table 4.**
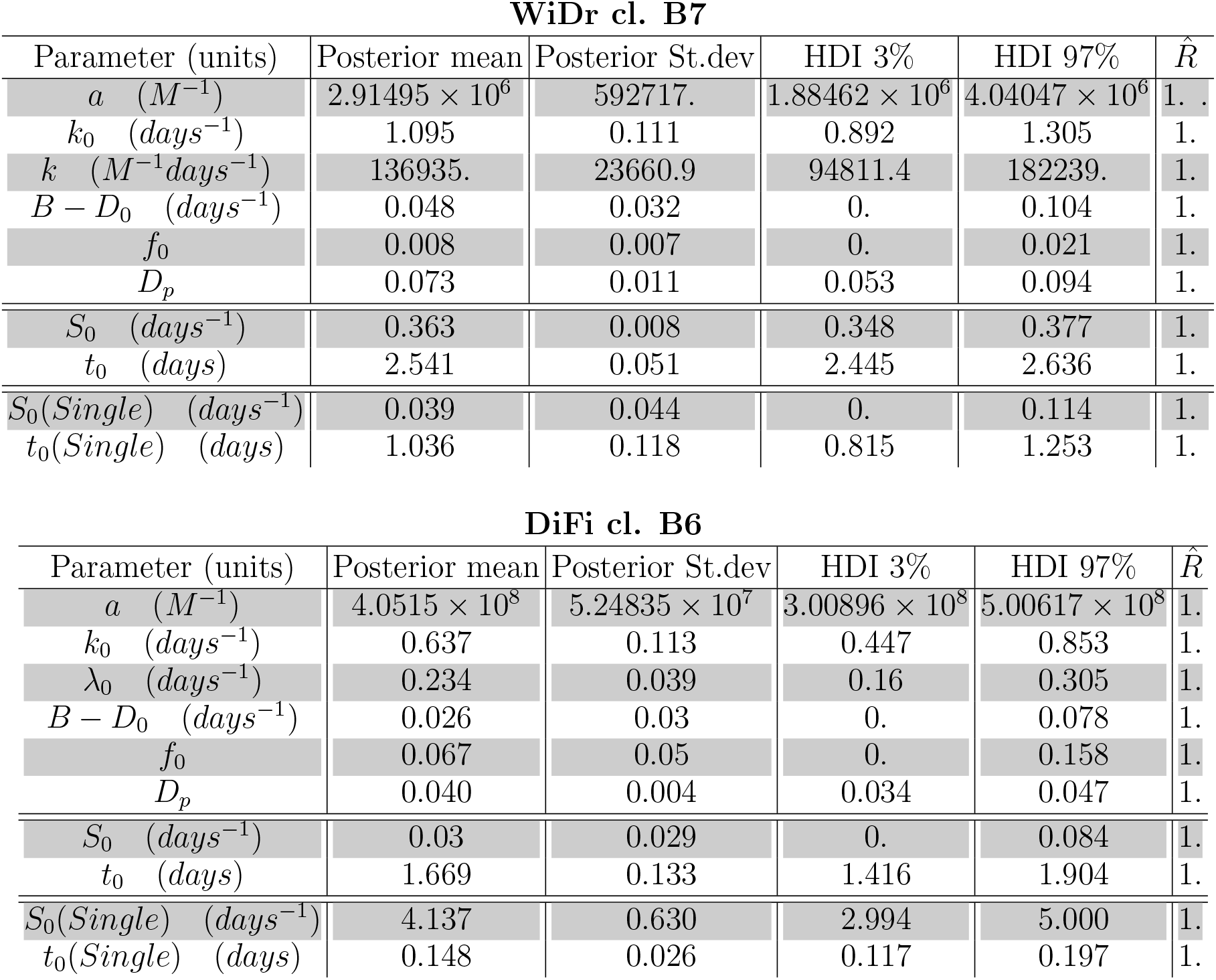
TP-model fit parameters. Tables list the values of the parameters of the TP model variant chosen (see methods). For each CRC clone, the posteriors distribution of the best model fit is reported including Mean value, Standard Deviation, Highest Density Interval (HDI) at 2.5%, Highest Density Interval (HDI) at 97.5% and the Gelman and Rubin’s convergence diagnostic statistics (*R*^). Values of *R*^ close to one indicate convergence to the underlying distribution. The model parameters are the following: ***a:*** inverse of the characteristic drug concentration. For a drug concentration *[M]<<a^-1^* cells die with their unperturbed death rate *D_0_*, while for concentrations *[M]>>a^-1^* the death rate ***κ*** reaches the maximal value (*D_0_ + k_0_*). ***k_0_*:** maximum death rate due to the drug. ***k*** coefficient of proportionality between drug concentration and rate of transition of sensitive cells to persister cells (λ=*k_0_[M]*). ***t_0_:*** time of the delay of the drug effect after administration. ***B-D_0_:*** growth rate in absence of drug (birthrate (*B*) minus death rate *D_0_*). ***S_0_:*** growth rate observed in the temporal window between drug administration and drug effect. ***λ_0_:*** rate of transition of sensitive cells to persister state. ***Dp:*** death rate of persister cells. ***t_0 (single):_*** delay in time of drug effect after treatment administration in the *single-dose assay. **S_0(single)_***: observed growth rate in the time window between treatment administration and drug effect (t < ***t_0(single)_***) in the *single-dose assay*.

**Extended Data Table 5.** Mutation rates of sensitive and persister cells according to experimental data using the MC-LD estimators. The table lists chronological mutation rates experimentally measured using the MC-LD fluctuation test. In the corresponding Fig. 4b we show the mean and SD of the values reported here for sensitive and persisters cells, for each cell line.

|  | WiDr cl. B7 |  | DiFi cl. B6 |  |
| --- | --- | --- | --- | --- |
| | $\hat{\mu}$ | $\hat{\sigma}$ | $\hat{\mu}$ | $\hat{\sigma}$ |
| Untreated Sensitive ($\times 10^{-7} \text{ days}^{-1}$) | 1.4 | 0.4 | 4.7 | 0.7 |
| Persisters ($\times 10^{-6} \text{ days}^{-1}$) | 7.0 | 0.4 | 3.0 | 0.3 |

## Notes

### Competing Interest Statement

A.Bardelli reports receiving commercial research grants from Neophore; is an advisory board member for Roche, Illumina, Guardant, and Third Rock; holds ownership interest in Neophore and Phoremost; and is an advisory board member for Horizon Discovery, Biocartis, and Neophore. All the other authors declare no competing interests.

